# Fungal Hyphae as Distributed Vapor Sinks

**DOI:** 10.64898/2026.05.13.724476

**Authors:** Yan Jun Lin, Leyun Feng, Awais Khan, Kyoo-chul Park, Sunghwan Jung

## Abstract

Hygroscopic surfaces act as local vapor sinks that reshape the condensation field around them, but whether distributed biological structures do the same has not been investigated. We have established that hyphae of fungal colonies functionally behave as vapor sinks, creating a dry region of width *δ* around themselves when placed on a cooled substrate. In addition, the radial distribution of droplet sizes steepens during condensation, and the rate at which droplets evaporate locally after chamber drying increases. In order to quantify this behavior, we employed a combination of time-resolved imaging and survival analysis to determine how long individual droplets persist on the surface surrounding the colony. These data were used to derive three quantitative measures of the vapor-sink effect. Each measure was found to be directly proportional to the vapor-sink strength of the substrate, as calibrated against NaCl–agar hydrogels of known water activity (LOOCV RMSE = 0.031 for recovered *a*_*w*_). These findings were consistent across three fungal genera (35 experiments), and all species fell along calibration lines defined by the hydrogel standards. This result is consistent with a diffusion-limited vapor-depletion framework. The measured genus-level *δ* ratios agreed to within 6% of predictions from structural absorbing capacity, and field measurements on *Gymnosporangium*-infected apple leaves were consistent with the same signatures under natural conditions. These results establish a non-contact method for inferring the material properties of thin hygroscopic biological surfaces from their condensation patterns.

---

Fungi require moisture, yet an excess will inhibit growth. Bacteria colonize the hyphae when the surface remains wet [1, 2], whereas germination and growth cease when the surface dries out. The cell wall mediates this interface. It is primarily composed of chitin, chitosan, and glucan, which together manage osmotic equilibrium, morphogenesis, and environmental sensing [3, 4, 5]. Additionally, the wall contains acetamide and hydroxyl groups that hydrogen-bond cooperatively [6], and the chitin microfibrils are organized into a porous scaffold with a high surface-area-to-volume ratio. Thus, the necessary components to support vapor uptake are present. Composites based on mycelia and chitosan films have demonstrated the capacity to sorb water under controlled conditions [7, 8]. However, no one has studied whether a living colony absorbs sufficient vapor to alter the condensation field surrounding it.

The main difficulty is resolving the vapor environment around a thin biological surface without damaging it; most current methods also lack the appropriate spatial resolution. Our hypothesis is that the hyphae operate as vapor sinks, with hydrogen bonding at the molecular scale and porosity at the network scale working together to enable uptake of atmospheric water vapor. To test this, we adopted a framework originally developed for abiotic hygroscopic sinks. In that framework, a soluble salt nucleus decreases local water activity, establishing a radial humidity gradient that prevents nucleation within a characteristic dry zone of width *δ* and slows droplet growth beyond it [9, 10]. Since then, this model has been refined for isolated hygroscopic droplets and solid objects [10, 11, 12, 13, 14, 15, 16], and is grounded in the physics of breath figures [17, 18, 19, 20, 21, 22, 23]. Engineered arrays of hygroscopic sinks are governed by the same diffusion-limited principles regardless of whether they consist of single points or competing arrangements [11, 12, 14, 24, 25]. However, none of these prior studies have examined distributed biological structures within this framework.

Here we image intact colonies of three fungal genera—*Aspergillus, Mucor*, and *Rhizopus*—through controlled condensation– evaporation cycles. We resolve each droplet in the surrounding vapor field individually and calibrate three condensation metrics against NaCl–agar hydrogel standards of known water activity.

## Results

### Diffusion-limited vapor-sink model

We treat the condensation pattern as an imprint of the quasi-steady vapor field above the cooled substrate. The water vapor concentration *c*(**x**) satisfies ∇^2^*c* = 0 (Laplace equation at fixed temperature), with far-field value *c*_∞_ and a boundary condition set by the vapor concentration at the boundary, linked to local water activity. A hygroscopic region with *a*_*w*_ < 1 imposes *c* ≈ *a*_*w*_ *c*_sat_(*T*_*s*_) at its surface, where *c*_sat_(*T*_*s*_) = *p*_*s*_(*T*_*s*_)*/R*_*g*_*T*_*s*_ is the saturation vapor concentration at the substrate temperature. The result is lateral gradients that shift where condensation and evaporation occur. Our experiments are in the diffusion-limited regime (*Da*_*s*_ ≳ 600 across plausible sticking coefficients; Methods), so the surface boundary condition reduces to its Dirichlet limit *c*(*R*_*s*_) = *a*_*w*_ *c*_sat_(*T*_*s*_).

For a sink of radius *R*_*s*_, we introduce a radial coordinate *ρ* measured from the sink center (converted below to *r* = *ρ* − *R*_*s*_, the experimentally measured distance from the sink edge, which is used throughout all figures and metrics), so that the sink surface is at *ρ* = *R*_*s*_ and the vapor-field domain is *ρ* ≥ *R*_*s*_. Solving ∇^2^*c* = 0 in spherical symmetry with far-field value *c*_∞_ and the Dirichlet boundary condition *c*(*R*_*s*_) = *a*_*w*_ *c*_sat_ (the diffusion-limited surface condition introduced above) gives

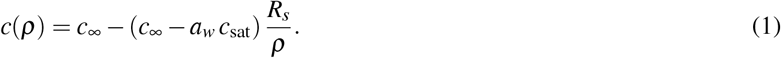

Defining the normalized vapor deficit *ϕ* (*ρ*) ≡ (*c*_∞_ − *c*(*ρ*))/(*c*_∞_ − *c*_sat_) and substituting Eq. 1 yields

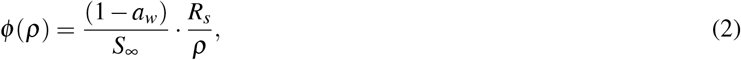

where *S*_∞_ = (*c*_∞_ − *c*_sat_) / *c*_sat_ is the far-field supersaturation. The key prediction, linear scaling of each observable with (1 − *a*_*w*_) at fixed chamber conditions, is verified experimentally in Fig. 2F.

The dry-zone boundary is defined by the condition *ϕ* > *ϕ*_crit_: wherever the local vapor deficit exceeds a critical threshold, supersaturation falls below the value *S*_crit_ required for heterogeneous nucleation and no droplets form. Setting *ϕ* (*ρ*_*δ*_) = *ϕ*_crit_ at the dry-zone edge *ρ*_*δ*_ = *R*_*s*_ + *δ* and solving Eq. 2 gives

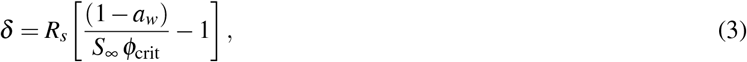

linear in (1 − *a*_*w*_) once chamber conditions (*S*_∞_, *S*_crit_, *R*_*s*_) are fixed. The fitted slope absorbs *ϕ*_crit_ and the geometric prefactors specific to our disk-on-plane setup; we standardized chamber conditions across every trial (Methods). So at fixed chamber conditions, any condensation observable that depends on *ϕ* should scale linearly with (1 − *a*_*w*_). We test it on three observables, dry-zone width *δ*, normalized droplet-size gradient, and half-attenuation distance *d*^*^, with operational definitions below and in Methods. The water activities we recover are effective, time-averaged values tied to NaCl–agar standards; they should not be read as instantaneous equilibrium values.

### Spatially organized condensation dynamics

It’s worth considering the raw experimental data prior to examining the metrics derived from them (Fig. 1). Fig. 1A (top row) shows an *Aspergillus* colony on the left and a NaCl–agar hydrogel (*a*_*w*_ = 0.75) on the right, both on the cooled substrate under brightfield illumination. Around each one there is a visible gap in the condensation. The middle and lower rows show close-up views at four representative timepoints: *t*_1_ = 10 min and *t*_2_ = 15 min during the condensation phase, and *t*_3_ ≈ 17 min and *t*_4_ ≈ 19 min during the evaporation phase (all measured from condensation onset). Each detected droplet is individually color-labeled. Droplets grow and coalesce throughout the condensation period (*t*_1_ to *t*_2_) except in the immediate vicinity of the sink, where growth suppression keeps them too small to merge. Following *t*_2_, the chamber begins to dehydrate. Drying begins nearest to the sink and progresses outward over the next few minutes (*t*_3_, *t*_4_).

**Figure 1.**
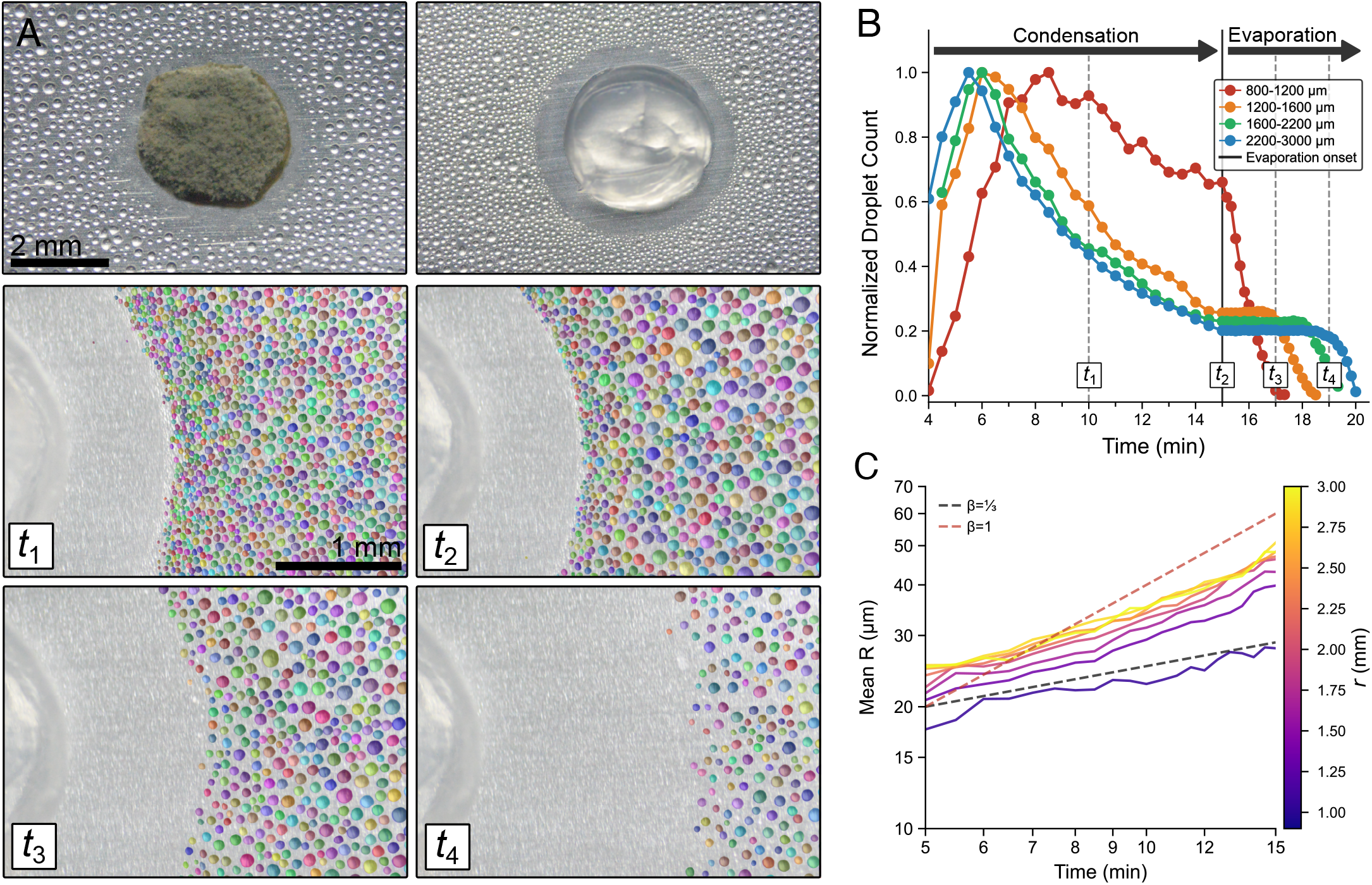
Time-resolved condensation dynamics around hygroscopic sinks. **(A)** Top row: an *Aspergillus* colony (left) and a NaCl–agar hydrogel with water activity (*a*_*w*_ = 0.75, right), each surrounded by a visible depletion zone (scale bar: 2 mm). Middle and bottom rows: close-up views at four timepoints (*t*_1_ through *t*_4_) showing Cellpose-segmented droplet overlays (scale bar: 1 mm). **(B)** Normalized droplet count (fraction of peak) versus time for distance bins from the source edge, 2:1 NaCl hydrogel trial (*a*_*w*_ = 0.75). Blue indicates far-field bins, which lose droplet count faster due to coalescence; red indicates near-field bins, which retain more droplets. Vertical line indicates chamber drying onset. **(C)** Droplet growth curves (mean radius versus time, log–log axes; same 2:1 NaCl hydrogel trial as panel B), binned by radial distance *r* from the source boundary. Radial distance is indicated using a plasma colormap (navy = near, yellow = far; colorbar units in mm). Reference slopes: *R* ∼ *t*^1/3^ (black, diffusion-limited) and *R* ∼ *t*^1^ (red, coalescence-dominated).

Survival curves (Fig. 1B) provide quantitative support for this ordering. Counts decrease throughout condensation, primarily through coalescence. Far-field bins lose droplets first; however, near-field bins persist longer since the droplets there are smaller and require more time to coalesce. The reverse occurs once drying begins: near-field bins cross 50% survival several minutes before the far-field bins, yielding a monotonic survival-time gradient. Distance-binned growth curves (Fig. 1C) exhibit the same two-regime behavior observed in breath-figure experiments: droplets closer to the sink follow *R* ∝ *t*^1/3^ [18] for longer, whereas far-field droplets transition to coalescence-dominated *R* ∝ *t*^1^ growth at an earlier timepoint. The difference between these two sets of curves is quantified by the size-gradient metric. Growth curves for all seven conditions are in Supplementary Fig. 3.

### Three metrics quantify vapor-sink strength

To test whether the linearity prediction (Eq. 2) holds experimentally, we examined four NaCl–agar hydrogel standards spanning *a*_*w*_ = 0.75–1.00 at 10% total solids (Methods; Fig. 2A–C, Supplementary Movies 1–2). Normalized droplet-size profiles 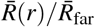 demonstrate that near-field suppression increases with increasing sink strength (Fig. 2E). Growth curves further confirm dynamic separation between near-field and far-field radii, where the former lags behind during condensation only when a sink is present (Fig. 2G).

**Figure 2.**
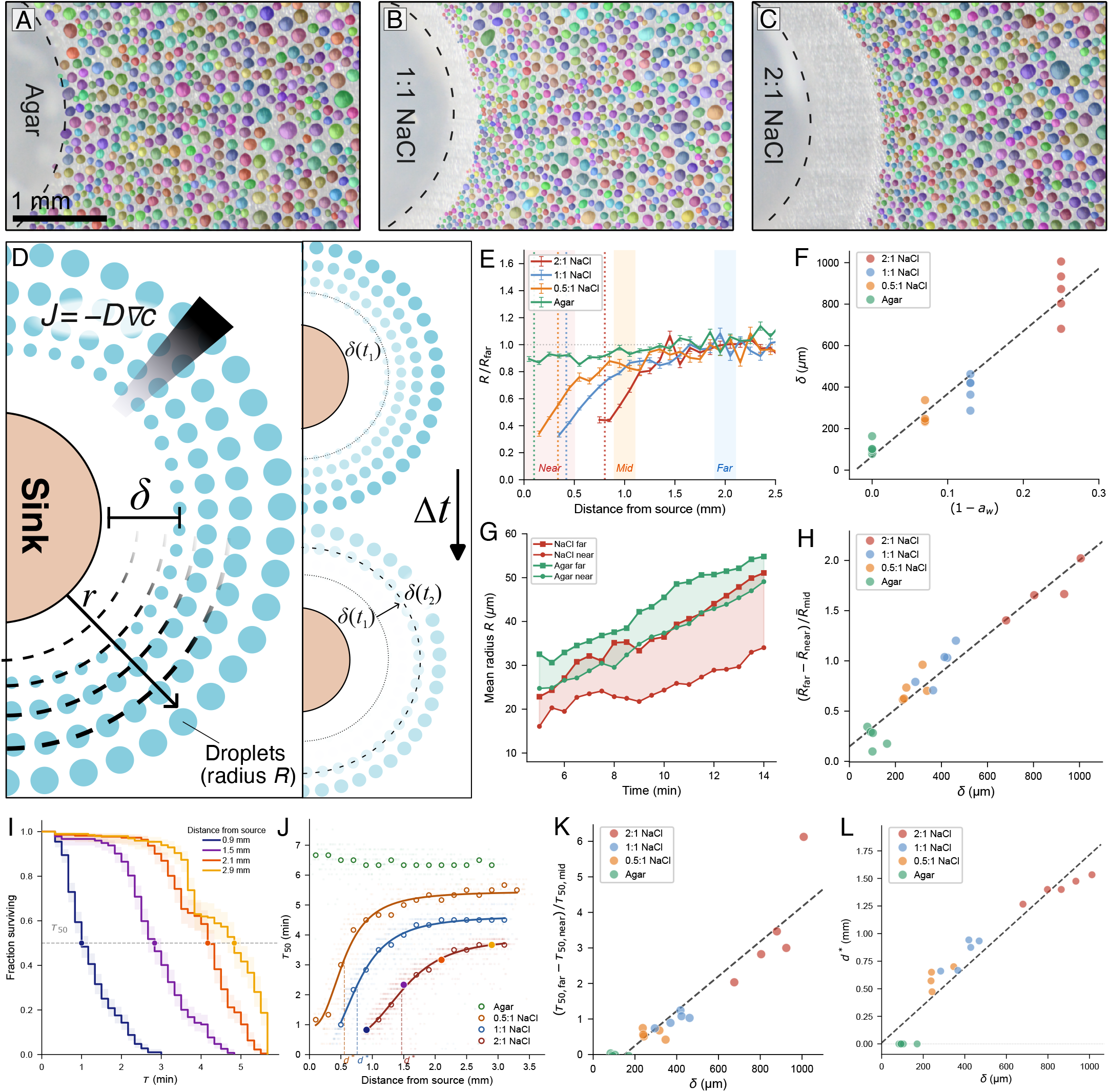
Three calibrated condensation metrics quantify vapor-sink strength. **(A–C)** Condensation micrographs with segmentation overlays for agar (*a*_*w*_ = 1.00), 1:1 NaCl (*a*_*w*_ = 0.87), and 2:1 NaCl (*a*_*w*_ = 0.75). **(D)** Schematic of dry-zone formation and expansion during evaporation. **(E)** 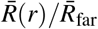 versus distance for all four hydrogel conditions (mean ± s.e.m.); shaded bands mark the near, mid, and far evaluation zones; dashed lines mark *δ* . Values exceeding 1.0 at large *r* reflect the use of a fixed-zone mean 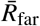 (1900–2100 µm) rather than the asymptotic far-field limit. **(F)** *δ* versus (1 − *a*_*w*_) for all 20 hydrogel trials. **(G)** Mean radius versus time for near- and far-field bins. **(H)** Normalized size gradient versus *δ* . **(I)** Kaplan–Meier survival curves at four distance bands pooled across all five 2:1 NaCl trials measured from evaporation onset; shaded bands: 95% CI; dots mark *τ*_50_ on the 0.5 survival line. **(J)** *τ*_50_ versus distance for four individual representative trials: agar, 0.5:1 NaCl, 1:1 NaCl, and 2:1 NaCl (circles: raw KM *τ*_50_; curves: Hill fits; dashed lines and italic labels: *d*^*^). **(K)** Normalized survival gradient (*τ*_50,far_ − *τ*_50,near_)*/τ*_50,mid_ versus *δ* (zone-based variant; see Methods). **(L)** *d*^*^ versus *δ* (*R*^2^ = 0.94, *n* = 20). Trials with flat *τ*_50_ profiles are assigned *d*^*^ = 0 by construction.

The dry-zone width *δ* exhibits a linear relationship with vapor-sink strength (1 − *a*_*w*_), and both the normalized droplet-size gradient and attenuation length collapse when plotted against *δ* (*R*^2^ ≥ 0.92, all *p* ≤ 3 × 10^−11^; Fig. 2F,H,L). Leave-one-out cross-validation of the *δ* calibration resulted in RMSE = 0.031 in water-activity units, demonstrating trial-to-trial reproducibility across all four hydrogel concentrations. Dry-zone width grew monotonically from the agar control to 2:1 NaCl (*δ* ≈ 860 µm), and the normalized droplet-size gradient covaried with *δ* across all 20 trials (Fig. 2F,H).

To investigate effects along the temporal axis, we applied Kaplan–Meier (KM) survival analysis (Fig. 2I–K). Distance-binned KM curves show forward lifetimes that decrease monotonically with proximity to the sink (Fig. 2I); the resulting *τ*_50_(*r*), the time at which 50% of droplets survive, profiles steepen with sink strength (Fig. 2J). A normalized survival gradient also scales linearly with *δ* (Fig. 2K), and the Hill-fit half-attenuation distance *d*^*^ scales linearly with *δ* (*R*^2^ = 0.94, *n* = 20; Fig. 2L), establishing that both the temporal and spatial aspects of nucleation suppression are governed by the same underlying vapor field. The normalized survival gradient is the temporal metric we apply to field specimens (Fig. 5F). Distance-stratified KM curves for all 20 hydrogel trials are in Supplementary Fig. 4.

### Universal collapse across fungal genera and hydrogel standards

The hydrogel calibration anchors each metric to (1 − *a*_*w*_). The next question is whether intact fungal colonies fall on the same axis. Three fungal genera were assayed: *Aspergillus* (dense conidial mold), *Mucor* (cottony aerial mycelium), and *Rhizopus* (sporangial mold), identified by macromorphology and light microscopy (Fig. 3A; Supplementary Movies 3–4).

The normalized droplet-size profiles 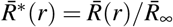 pooled over all 35 trials lay out a monotonic spectrum of vapor-sink strength that bridges biotic and abiotic sources, in the order 2:1 NaCl > 1:1 NaCl > 0.5:1 NaCl ≈ *Aspergillus > Mucor* ≈ *Rhizopus >* agar (Fig. 3B). The per-trial heatmap (Fig. 3C) shows that within-group variability is modest relative to between-group separation. Individual trial profiles are in Supplementary Fig. 2. All 35 trials, biotic and abiotic, fall on a single *d*^*^–*δ* trend (*R*^2^ = 0.90, *n* = 35; Fig. 3D) and a single droplet-size-gradient–*δ* trend (*R*^2^ = 0.91, *n* = 35; Fig. 3E). The size-gradient collapse is robust across 36 zone-boundary combinations (*R*^2^ = 0.83–0.93; Supplementary Table 1). Of the three primary observables, only *δ* separated the three genera by Kruskal–Wallis test (*p* < 0.05); *d*^*^ also separated genera (*p* = 0.048).

*Aspergillus* is the strongest of the three sinks (*δ* ≈ 300 µm, comparable to the 0.5:1 NaCl hydrogel at *a*_*w*_ = 0.93); *Mucor* and *Rhizopus* produce weaker depletion (*δ* ≈ 110–140 µm). The architectural basis for the *Aspergillus* excess is examined next, using the two genera that established mature aerial colonies under the standardized growth conditions.

**Figure 3.**
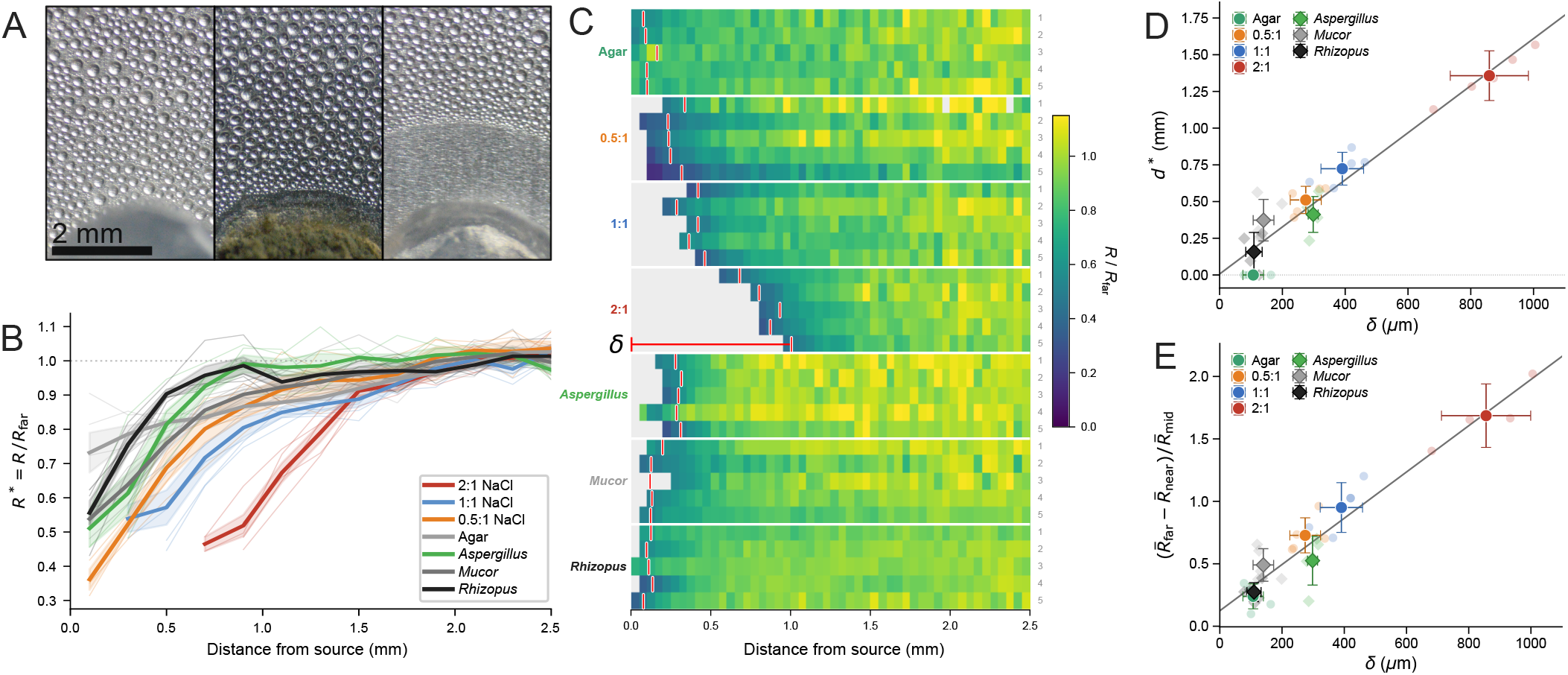
Vapor-sink metrics collapse across fungi and hydrogels. **(A)** Condensation micrographs around an agar control (left), *Aspergillus* colony (center), and 2:1 NaCl hydrogel (right). Scale bar: 2 mm. **(B)** Normalized radial profiles 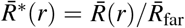 for all 35 trials (thin lines) and group means ± SEM (thick lines). 200 µm bins; *t* = 14.5–15.5 min. **(C)** Heatmap of 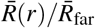 for all 35 trials, grouped by condition. Gray: dry zone; red dashed ticks: *δ* . **(D)** Half-attenuation distance *d*^*^ versus dry-zone width *δ* for all 35 trials (circles: hydrogel trials; diamonds: fungal trials). **(E)** Normalized droplet-size gradient 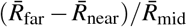 versus *δ* (*R*^2^ = 0.91, *n* = 35).

### Colony architecture accounts for vapor-sink strength

To pin down why *Aspergillus* is the stronger sink, we characterized colony morphology in the two genera that produced viable colonies — *Aspergillus* and *Mucor* — with four metrics that reduce to three axes: volumetric hyphal load (structure thickness, erosion survival), surface roughness (FFT spectral slope), and intra-hyphal heterogeneity (Hessian tubeness CV) (Fig. 4). The dense conidia-dominated surface of *Aspergillus* contrasts qualitatively with the hypha-dominated surface of *Mucor* (Fig. 4A,B; *Rhizopus* excluded for insufficient aerial mass); four quantitative metrics substantiate this.

**Figure 4.**
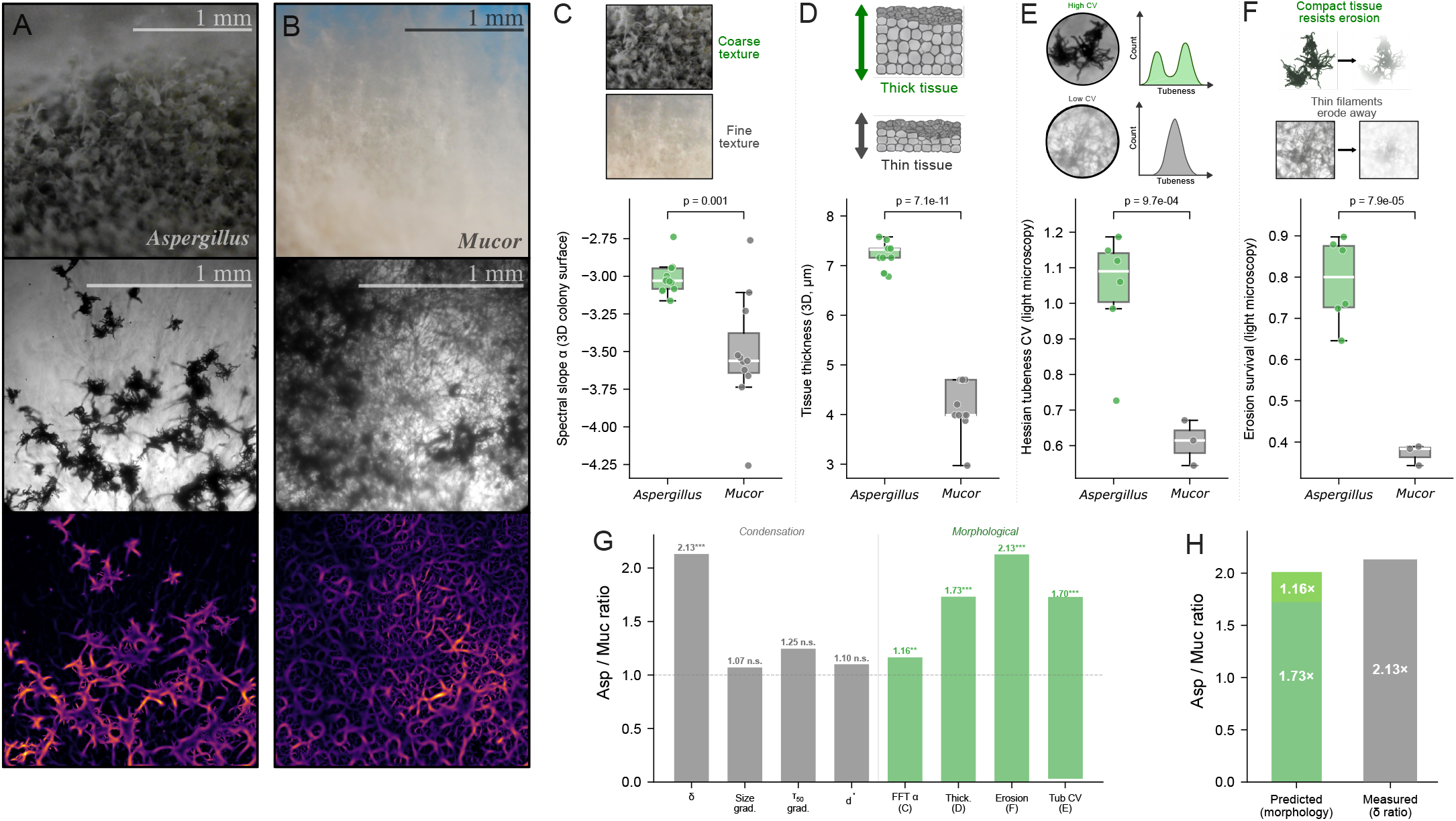
Colony architecture accounts for vapor-sink strength. **(A, B)** Three-channel colony imaging of *Aspergillus* (A) and *Mucor* (B): color macro brightfield (top), grayscale light microscopy (middle), and local-density heatmap (bottom). Scale bars: 1 mm. **(C)** FFT spectral slope *α* from macro-scale brightfield (Welch’s, *p* = 0.001); shallower = rougher. **(D)** Median structure thickness from distance-transform of colony-surface masks (Welch’s, *p* = 7.1 × 10^−11^). **(E)** Hessian tubeness CV (Welch’s, *p* = 9.7 × 10^−4^; MWU bounded by *p* ≥ 0.024 at *n*_*a*_ = 6, *n*_*m*_ = 3); high CV indicates bimodal solid-core/filamentous-edge hyphae. **(F)** Erosion survival fraction at iteration 10 (Welch’s, *p* = 7.9 × 10^−5^; MWU bounded by *p* ≥ 0.024). **(G)** Asp/Muc ratios for four condensation observables (gray) and four morphological metrics (green); all eight exceed 1 (sign test *p* = 0.008). Ratios are oriented *Aspergillus*-favored; for *α* this means |*α*_Muc_ | / | *α*_Asp_ | (both slopes negative). ^*^ *p <* 0.05, ^**^ *p <* 0.01, ^***^ *p <* 0.001. **(H)** Absorbing-capacity decomposition: thickness (1.73 ×) × hyphal fraction (1.16 ×) predicts a *δ* ratio of 2.01, within 6% of the measured 2.13 ×. Boxes (C–F): IQR; line: median; whiskers: 1.5 × IQR; dots: ROIs (*n* = 13/11 colony-surface; *n* = 6/3 light-microscopy).

Four metrics from the established image-analysis literature [26, 27, 28, 29] distinguish the two architectures (Fig. 4C– F; per-metric means, Welch’s *t*, and Cohen’s *d* in Supplementary Table 2). *Aspergillus* has a shallower FFT spectral slope (richer fine-scale structure), 1.73 × greater median structure thickness, 1.70 × greater heterogeneity in Hessian tubeness CV (a bimodal fingerprint of mixed solid cores and filamentous edges), and 2.13 × more compact hyphae (higher erosion survival). All four metrics survive independent robustness checks against magnification, segmentation threshold, and an orthogonal GLCM-contrast cross-validation (Supplementary Figs. 9, 12, 14). The four reduce to three independent axes: volumetric hyphal load (thickness, erosion survival), surface roughness (FFT slope), and intra-hyphal heterogeneity (tubeness CV).

We define the *absorbing capacity* 𝒜 = *f*_tissue_ × *d*_structure_, the product of hyphal area fraction *f*_tissue_ and median structure thickness. By the geometric identity *V*/*A*_proj_ = *f*_tissue_ · *d*_structure_, 𝒜 is total hygroscopic hyphal volume per unit projected area, the image-derived analogue of mass per footprint (Supplementary Fig. 13). Because *Aspergillus* and *Mucor* share similar chitin– glucan wall composition and bulk sorption capacity [3, 30, 8, 7], the volume ratio is the relevant comparison. Decomposing the *Aspergillus*/*Mucor* ratio gives 1.73 (thickness) × 1.16 (hyphal area fraction *f*_tissue_, computed from the same binarized colony-surface masks; see Methods) = 2.01, within 6% of the measured *δ* ratio of 2.13 (Fig. 4H). With *n* = 2 genera the agreement is suggestive rather than definitive, but the magnitude and direction match.

The perimeter-to-area ratio (image-derived specific surface area) is 1.78 × higher in *Mucor* (Supplementary Fig. 13D), consistent with its thinner filamentous structures. *Aspergillus* nonetheless produces the stronger sink, which rules out interfacial area as the operative quantity (further discussion of internal equilibration timescales in Supplementary Text 1). Total absorbing mass per projected area sets sink strength: *Aspergillus*’s larger reservoir keeps its time-averaged surface water activity below *Mucor*’s across the 15-min window, matching the measured ∼ 2 × ratio in *δ* . All eight *Aspergillus*-vs-*Mucor* ratios point in the same direction (sign test *p* = 0.008; Fig. 4G).

### Field validation on *Gymnosporangium*

To test whether the laboratory signatures extend to natural pathosystems, we imaged apple leaves colonized by *Gymnosporangium* (cedar-apple rust), which develop clustered aecia that serve as localized sources of aeciospores, under the same standardized protocol (*N* = 6: *n* = 3 healthy, *n* = 3 diseased with secondary saprophytic colonization; Fig. 5). These fungi cannot be grown in laboratory culture.

Representative micrographs (Fig. 5A) show the condensation field surrounding healthy (left) and diseased (right) specimens. In both cases, droplets near the fungal hyphae are visibly smaller and sparser than those at greater distances, consistent with the laboratory observations. Manual expert annotation of individual droplets within a defined analysis region (Fig. 5B) yielded a significant positive correlation between equivalent radius and distance in all six trials (individual trial Pearson *r* = 0.43–0.66, all *p <* 10^−6^; *N* = 1,507 total droplets; sign test for all-positive slopes: two-tailed *p* = 0.031). The slope of the mean droplet-size as a function of distance was 10.5 ± 3.0 µm mm^−1^ (mean ±s.d.), and far-field droplets were on average 2.3 ± 0.6-fold larger than near-field droplets. Individual trial scatter plots are shown in Supplementary Fig. 8.

**Figure 5.**
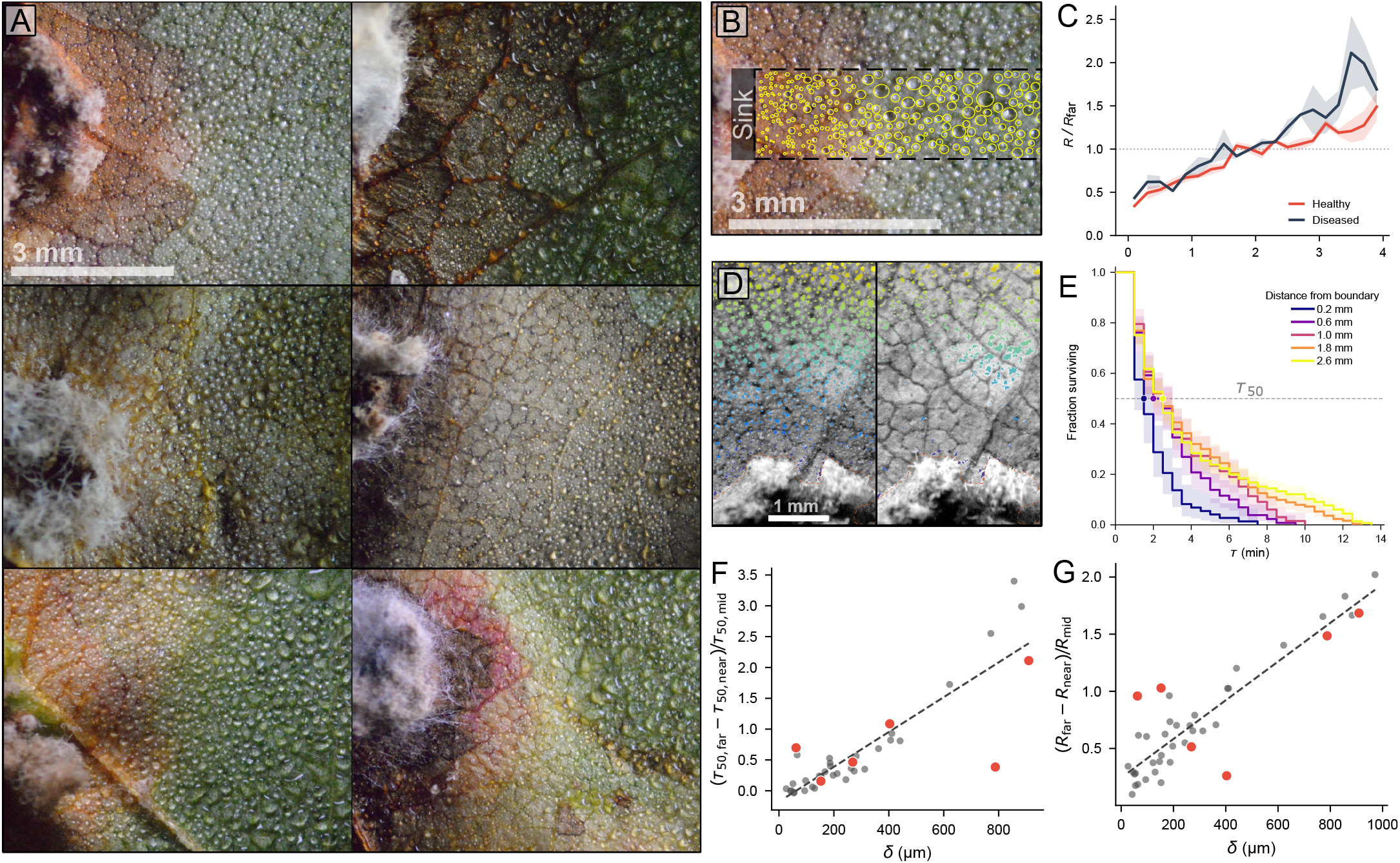
Vapor-sink signatures in natural fungal pathosystems. **(A)** Condensation micrographs on *Gymnosporangium*-infected apple leaves. Left: healthy rust specimens; right: diseased specimens with secondary fungal colonization. Scale bar: 3 mm. **(B)** Example of manual droplet annotation; yellow circles mark individual droplets used to compute the size–distance relationship. Scale bar: 3 mm. **(C)** Normalized radial profiles 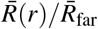 for healthy and diseased specimens (mean ± s.e.m.). **(D)** Example of ilastik pixel classification of a healthy rust specimen (left: evaporation onset; right: ∼ 75% complete), used to quantify droplet evaporation gradient where individual droplet tracking is not feasible due to low-contrast field. Scale bar: 1 mm. **(E)** Kaplan–Meier survival curves at five distance bands of a single trial measured from evaporation onset; shaded bands: 95% CI. **(F)** Normalized survival gradient (*τ*_50,far_ − *τ*_50,near_)*/τ*_50,mid_ versus *δ* for field specimens (red) overlaid on calibration data (gray). **(G)** Normalized size gradient versus *δ* for field specimens overlaid on laboratory data.

Normalized radial profiles 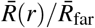 (Fig. 5C) confirm that both healthy and diseased specimens exhibit the characteristic monotonic increase in droplet size with distance from the fungal boundary. The two groups largely overlap, indicating comparable vapor-sink strength despite the difference in hyphal volume (*t* = − 0.79, *p* = 0.47; *n* = 3 per group).

Droplet density was quantified by ilastik pixel classification (Fig. 5D). Distance-binned KM curves (Fig. 5E) show that near-field bins evaporate first, with the front propagating outward (Supplementary Movie 5), reproducing the spatiotemporal ordering observed in laboratory trials (Fig. 1B). Overlaid on laboratory calibration, field specimens fall within the size-gradient–*δ* trend (Fig. 5G) and on a zone-based survival-gradient variant of the temporal metric (Fig. 5F; Methods).

## Discussion

Three major conclusions emerge from this study. First, all three phyla of fungal hyphae examined here act as vapor sinks: Ascomycota (*Aspergillus*), Mucoromycota (*Mucor, Rhizopus*), and a field-collected Basidiomycete (*Gymnosporangium*). Second, the three calibrated metrics (*δ*, the normalized droplet-size gradient, and *d*^*^) show that biotic and abiotic sources collapse onto the same scaling curves and yield an effective water activity at LOOCV RMSE = 0.031. Third, genus-level differences in *δ* correlate with colony architecture (absorbing capacity, mass per projected area), not wall chemistry or specific surface area. Chitin-rich walls are recognized as being able to sorb large amounts of water [7, 8], and it has been shown that fungal evaporative cooling can produce incidental condensation [31]; however, no previous studies have treated intact fungi as vapor sinks in their own right.

Previous studies examining hygroscopic sinks relied on endpoint imaging and manual analysis [10, 11, 12]. Rather than capturing images at endpoints and analyzing them manually, our approach captures entire condensation–evaporation cycles using deep-learning segmentation [32] and KM survival analysis on tens of thousands of individually tracked droplets per trial, which provides the ability to identify temporal dynamics that endpoint methods cannot detect.

There are several constraints on the framework. Our planar patches operate at *R*_*s*_*/δ* ≈ 1.7–14, in the boundary-layer regime where depletion is dependent on the thermodynamic boundary condition of the sink rather than its geometry; this eliminates the dependence of chemistry on morphology and explains why fungi and hydrogels collapse onto shared metric–metric relationships. Linearity assumes fixed far-field supersaturation, isolated sinks, and *Da*_*s*_ ≫ 1. The *Gymnosporangium* aecia (millimeter-scale, well-separated) satisfy these conditions. Extending the framework to continuous mycelial mats, where inter-hyphal spacing falls below *δ* and depletion zones begin to overlap, will require models that include multiple sinks competing for vapor [11]. A controlled experiment is difficult to perform with biological samples because identical patches cannot be reliably reproduced; arrays of artificial hygroscopic disks at controlled spacing offer a practical way to determine whether the calibration established here applies to dense, continuous coverage.

The Asp/Muc ordering in *δ* is surprising on the basis of chemistry alone. *Mucor*’s chitin–chitosan-rich walls [33] should adsorb more water per unit mass than *Aspergillus*’s glucan-rich walls [3, 30]. However, *Aspergillus* creates larger depletion zones, indicating that architecture rather than chemistry determines *δ* . Whether the RodA hydrophobin rodlet layer on *Aspergillus* influences vapor uptake at the molecular scale is unclear at this time [31, 34] and will need to be addressed in future work. The predicted absorbing-capacity value (2.01 ×, within 6% of the measured *δ* ratio of 2.13; Fig. 4H) supports this conclusion, though *n* = 2 genera limits the predictive capability of the model. Quantitative separation of intrinsic cellular water activity from architectural amplification will require independent dynamic vapor sorption measurements on milligram-scale colony samples, a measurement that would necessarily destroy the intact architecture whose collective effect is studied here.

The structural and thermodynamic perspectives can be connected through capillary condensation. Approximating inter-hyphal spaces as pores of effective radius *r*_*p*_, the Kelvin equation gives

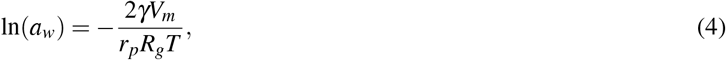

where *γ* is surface tension, *V*_*m*_ the molar volume of water, *R*_*g*_ the gas constant, and *T* temperature. To first order (1 − *a*_*w*_) ∝ 1/*r*_*p*_, so from Fig. 2F, *δ* ∝ 1/*r*_*p*_: the measured *δ* ratio of 2.13 implies that *r*_*p*_ in *Aspergillus* is roughly half that of *Mucor*. If both colonies capture comparable vapor volumes, fractional pore saturation scales inversely with total absorbing capacity 𝒜, and the thermodynamics-based *δ* ratio (2.13) converges on the structure-based *A* ratio (2.01), indicating that hyphal volume dictates the capillary dynamics governing the vapor sink.

It has previously been shown that fungi can cool themselves by evaporation, acting as vapor *sources* that produce condensation on surfaces above the colony [31]. The mechanism described here is the converse: the colony edge acts as a vapor *sink*, absorbing water vapor laterally and depressing local supersaturation. An inert NaCl–agar hydrogel held at the same temperature reproduces *δ*, confirming that osmotic vapor-pressure depression, not thermal depression, is sufficient to explain the effect. Both effects can coexist but operate on different physical axes and should not be conflated. Vapor-sink activity is nonetheless a form of microhabitat engineering at the atmospheric boundary layer. In substrate systems, fungi are known to create their own microenvironment. For example, wood-decay fungi regulate moisture levels in colonized wood as a competitive strategy, controlling internal hyphal water transport to rehydrate distant microhabitats in otherwise dry environments [35, 36, 37]. Our result extends this principle from substrate liquid water to atmospheric water vapor: the colony passively reshapes the vapor boundary layer within *δ* of its edge through the collective hygroscopicity of the chitin–glucan wall, and the magnitude of this environmental modification is proportional to a measurable structural parameter (absorbing capacity, Fig. 4H).

Bacteria travel through continuous liquid films on hyphal surfaces, the “fungal highways” described for pollutant-degrading bacteria [38] and mycosphere migrators [39]. Biofilm formation on *Aspergillus niger* hyphae is dependent upon secreted matrix components that require aqueous conditions [40]. On leaf and soil surfaces, bacteria survive daytime desiccation only within microscopic surface wetness: sub-micrometer liquid films maintained by deliquescence of hygroscopic particles, which are typically invisible to conventional leaf-wetness sensors [41, 42]. The dry-zone widths measured here (*δ* ≈ 100–860 µm) exist at the spatial scale at which these films determine microbial survival. If vapor-sink activity were to thin or eliminate near-field water films, it could provide a physical mechanism for limiting the bacterial colonization of hyphal surfaces noted in the introduction [1, 2].

A similar relationship applies to plant disease. The majority of foliar fungal pathogens require sustained leaf wetness for spore germination and infection [43], and forecasting models parameterize infection risk as a function of temperature and leaf-wetness duration [44]. However, leaf-wetness duration remains a non-standardized meteorological variable, and bulk humidity proxies fail to capture the microscale water films that determine infection success [45]. A number of pathogens can actively reshape their local microclimate to facilitate infection: bacterial effectors establish aqueous apoplastic environments under high humidity [46], and rust aeciospore liberation depends on water perfusion of inter-spore gaps at the millimeter scale [47]. The dry-zone widths reported here (*δ* ≈ 100–860 µm) fall within the range of sizes relevant to both the surface water films required for germination and the aecial structures from which our *Gymnosporangium* field signatures arise. Whether a fungal lesion’s vapor-sink activity locally suppresses or augments these films, and how the resulting boundary-layer microclimate affects infection efficiency, sporulation, and epidemic development, are testable questions. The condensation-pattern metrics developed here may offer a non-contact readout of leaf-surface humidity near fungal lesions, complementing the existing imaging toolkit of RGB, hyperspectral, and thermal pre-symptomatic detection [48] with a biophysically interpretable measurement that is not otherwise available.

At the single-spore scale (∼ 10 µm), basidiospore ballistospory is powered by hygroscopic sugars: mannitol and hexoses secreted at the hilar appendix condense Buller’s drop from saturation vapor in the ambient air, and coalescence of this drop with the adaxial drop accelerates the spore to > 10^4^ *g* [49, 50]. *Detached basidiospores re-wet in humid air and act as effective cloud condensation nuclei [51]*. *At the conidial scale, Penicillium rubens* conidia formed at low water activity accumulate compatible solutes (glycerol) and attract significantly more water from humid air than conidia formed at high *a*_*w*_, as measured by dynamic vapor sorption [52]. At the colony scale, the chitin–glucan polymer network absorbs water vapor across the mycelial surface in such a way as to generate a measurable dry zone. The shared physics across all three scales is vapor-pressure depression by hygroscopic biological materials: sugars for spores, compatible solutes for conidia, structural wall polymers for colonies. Recovery of the same signature across representatives of three fungal phyla diverged by hundreds of millions of years (Ascomycota, Mucoromycota, Basidiomycota) is suggestive of a general physical consequence of exposing a chitin-rich surface to humid air rather than a lineage-specific adaptation, though broader taxonomic sampling will be necessary to settle the point. The absorbing-capacity framework (Fig. 4H) yields the colony-scale quantification: here too, sink strength scales with total polymer mass per projected area. Whether this framework predicts *δ* across a wider taxonomic range is a natural follow-up experiment.

The temporal metrics have one methodological caveat. During evaporation, near-field droplets are on average smaller because their growth was suppressed during condensation, and smaller droplets evaporate intrinsically faster due to their higher curvature and larger surface-to-volume ratio (the Kelvin effect on droplet vapor pressure), regardless of any ongoing vapor depletion by the sink. The half-attenuation distance *d*^*^ and the normalized survival gradient therefore combine two contributions: (i) size-dependent intrinsic evaporation rates inherited from the condensation phase, and (ii) any persistent vapor uptake by the sink during evaporation. Because the same convolution is present in the hydrogel calibration trials, the temporal metrics still collapse with *δ* on a single scaling relationship (Fig. 2K,L; Fig. 3D), and the resulting calibration consistently captures the joint signature across both biotic and abiotic sinks, albeit without separating the two contributions. Untangling the two would require an independent measurement of surface water activity at evaporation onset, which is outside the present scope.

The field dataset is small (*N* = 6), limited by the obligately biotrophic life cycle of *Gymnosporangium*. Outside fungi, the same framework should be applicable to any thin hygroscopic surface where gravimetric measurement is impractical. Biofilms, pollen, insect cuticles, and synthetic atmospheric water-collection coatings [53, 54] are prime candidates. For engineered hygroscopic surfaces, our results afford a design insight: the filamentous networks examined here operate at *R*_*s*_*/δ* ≈ 1.7–14, in the thermodynamically constrained regime where depletion is controlled by boundary-condition chemistry rather than collector geometry [11, 14, 55]. In this regime, total sorbent mass per unit footprint, not surface-area-to-volume ratio, sets vapor-capture performance, a principle supported quantitatively by the *Mucor* result (higher specific surface area, weaker sink).

## Methods

### Experimental setup and sample preparation

Experiments were conducted in a sealed imaging chamber equipped with a Peltier-cooled aluminum substrate, a closed-loop water chiller (CW-3000), and time-lapse imaging (Supplementary Fig. 1). The thermoelectric module was thermally coupled to the substrate through thermal paste; silane-treated aluminum foil served as the cold-side surface (*T*_*s*_ ≈ 7 ^◦^C). Water vapor was supplied by a compressor nebulizer (InnoSpire Essence, Respironics) at constant output. Relative humidity was monitored by a digital hygrometer–thermometer (IPT-100S, Elitech) with external temperature probe bound to the Peltier surface, logging at 1-s resolution. Time-lapse imaging used a motorized focus rail (StackShot, Cognisys), LED illumination, and a DSLR camera in tethered mode (shutter 1/125 s, ISO 6400, DX format 3936 × 2624 px).

Fungal cultures were isolated by serial subculture on potato dextrose agar at 22 ± 2 ^◦^C until homogeneous colonies were obtained (2–3 transfers). Three recurring genera were identified by macromorphology and light microscopy based on characteristic reproductive structures: *Aspergillus* (dense, powdery-granular conidial mold with vesicle-bearing conidiophores), *Mucor* (cottony white aerial mycelium with columellate sporangia), and *Rhizopus* (darkly pigmented sporangial mold with rhizoids and stolons). Identification was at the genus level. Circular patches (3 mm diameter) were excised with a sterile disposable dermal punch (3 mm, Miltex) and transferred onto the cooled foil with aerial mycelium facing upward. NaCl–agar hydrogel disks (radius *a* = 1.5 mm, height *h* ≈ 500 µm; 10% total solids) were cast at three NaCl:agar mass ratios, 2:1 (*a*_*w*_ ≈ 0.75), 1:1 (*a*_*w*_ ≈ 0.87), and 0.5:1 (*a*_*w*_ ≈ 0.93), with water activities approximated from NaCl solution data at matched salt mass fraction [56]. The hydrogel *a*_*w*_ values above are nominal references, and we report all downstream results on a calibration-defined scale rather than against absolute *a*_*w*_. Agar-only disks of identical geometry served as geometric controls (*a*_*w*_ = 1.00).

Before each run, the chamber was sealed and equilibrated. The chiller, power supply, illumination, nebulizer, and camera were activated simultaneously. During condensation (0 to 15 min), humidity increased under constant nebulizer output while temperature and humidity were logged at 1-second resolution. For evaporation, the nebulizer and Peltier were switched off and imaging continued until all droplets disappeared. Temperature and humidity traces were reproducible across 13 trials (temperature pooled SD = 0.81 ^◦^C; humidity pooled SD = 1.79% RH across the full condensation window; Supplementary Fig. 15), which supports the assumption that chamber conditions were standardized across experiments.

### Image analysis

Droplet segmentation for laboratory cultures and hydrogel trials used Cellpose (v4.0.8) [32] with the pretrained cyto3 model (no fine-tuning). Inference parameters: `diameter` = 90 px, `flow threshold` = 0.2, `cellprob threshold` = 1.0. Morphometric properties (area, equivalent radius, centroid) were extracted via scikit-image `regionprops`.

Distance from each droplet centroid to the nearest point on the source boundary was computed using the Euclidean Distance Transform (EDT). For fungal colonies with irregular morphology, source boundaries were defined by freehand polygon tracing; for hydrogel disks, an analytic ellipse boundary was used. All distances are measured from the source (patch) edge. Three primary metrics were extracted from each trial:

#### Dry-zone width (*δ*)

The source boundary was sampled at 100 equally spaced points. For each sample, all droplets whose nearest boundary point was that sample were identified, and the minimum EDT distance was recorded. The dry-zone width is the mean of these per-ray minima over the late-condensation window (10–15 min). The raycast procedure and *δ* values for all 35 trials are shown in Supplementary Fig. 7.

#### Normalized size gradient

At the 15-minute timepoint (*t* = 14.5–15.5 min), droplets were binned in 100 µm distance increments from the source boundary and mean radius computed per bin. Bins containing fewer than 5 droplets were assigned a mean radius of zero. Three zones were defined: near (0–500 µm), mid (900–1100 µm), and far (1900–2100 µm), with zone means computed over all constituent bins. The metric 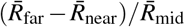 captures the magnitude of the radial size gradient normalized to the mid-field droplet scale. A sensitivity analysis over 36 alternative zone boundaries confirms robustness (*R*^2^ = 0.83–0.93; Supplementary Table 1).

#### Normalized survival gradient

The normalized survival gradient (*τ*_50,far_ − *τ*_50,near_) / *τ*_50,mid_ was computed from the same near (0–500 µm), mid (900–1100 µm), and far (1900–2100 µm) evaluation zones as the size gradient, using zone-mean Kaplan–Meier *τ*_50_ values in place of zone-mean radii. A sensitivity analysis over nine combinations of bin width and minimum track count confirms that *τ*_50_–distance profiles are insensitive to binning parameters (Supplementary Fig. 5).

#### Half-attenuation distance (*d*^*^)

Droplet lifetimes were defined as *τ*_fwd_ = *t*_death_ − 900 s (forward lifetime from evaporation onset at *t* = 15 min). Coalescence events were right-censored. Minimum track length: 3 frames. Droplets were binned in 200 µm distance increments (minimum 15 droplets per bin). For each bin, median survival time *τ*_50_ was estimated via Kaplan– Meier fitting (lifelines package). *d*^*^ was extracted by fitting a four-parameter saturating function *T* (*d*) = *T*_0_ + *Ad*^*n*^/(*d*^\**n*^ + *d*^*n*^) via bounded nonlinear least squares (SciPy curve fit; minimum 3 valid bins). Trials returning a flat *τ*_50_ profile (no detectable gradient) were assigned *d*^*^ = 0.

### Colony architecture analysis

Colony architecture was measured using four metrics for surface texture and tissue geometry, computed on hand-selected regions of interest (ROIs) from each colony image, using the same DSLR macro system as the time-lapse experiments (∼ 1 µm/px, calibrated with a stage micrometer). Macro-scale brightfield colony-surface photographs (single-plane, not focus-stacked) were binarized by adaptive thresholding into tissue and air phases. Analysis used custom Python tooling (NumPy, SciPy, scikit-image). Comparison of colony architecture was restricted to the two genera that established viable colonies under the standardized growth conditions: *Aspergillus* and *Mucor*. A total of 13 *Aspergillus* and 11 *Mucor* ROIs were used for macro-scale colony-surface metrics, and 6 *Aspergillus* and 3 *Mucor* images were used for light-microscopy metrics.

*FFT spectral slope α* (Fig. 4C): each macro-scale brightfield colony-surface photograph was tiled into 256 × 256 px windows with 128 px stride, Hann-windowed, 2D Fourier-transformed, and radially averaged. The radial power spectrum was fit on log–log axes over the band 0.02–0.40 cycles/px. A shallower (less negative) slope indicates a greater amount of surface roughness at the microscopic level. Spatial-domain GLCM contrast was also computed as a separate cross-validation metric (distance = 1 px, four orientations averaged) and confirmed the FFT genus ordering (Supplementary Fig. 9).

*Structure thickness* (Fig. 4D): the tissue phase of each binarized colony-surface mask was distance-transformed (Euclidean), and the median distance-transform value within the tissue was recorded as the structure thickness in µm. Sensitivity to thresholding was evaluated by sweeping the adaptive multiplier *k* = 0.3–0.7 × std; both genera fall within published hyphal-diameter ranges across the entire sweep, and the Asp/Muc ratio remains consistent at 1.67–1.73 (Supplementary Fig. 14).

*Hessian tubeness CV* (Fig. 4E): multi-scale Hessian tubeness response was computed on grayscale light microscopy at scales *σ* = 1, 2, 4, 8, 16 px and the maximum response across scales retained per pixel; representative tubeness response and tissue-mask overlays for each genus are shown in Supplementary Fig. 10. Within each adaptive-threshold tissue mask, the coefficient of variation 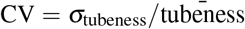 was computed. A high CV value implies a bimodal distribution within the tissue, consisting of solid core regions (low tubeness) and filamentous edge regions (high tubeness). Hessian tubeness CV was chosen from among other Hessian-derived metrics because it best represented biologically interpretable information and was independent of magnification; the Asp/Muc ratio is preserved across magnifications (1.71 × at 10 ×, 1.76 × at 20 ×, 1.65 × at 40 ×; Supplementary Fig. 12).

*Erosion survival* (Fig. 4F): the binarized light-microscopy mask was eroded iteratively using a disk structuring element (radius = 1 pixel per iteration) for ten iterations, and the fraction of tissue pixels remaining after ten iterations was recorded. Values closer to one indicate more compact and structurally resistant tissue. Examples of successive erosion iterations for each genus are provided in Supplementary Fig. 11.

*Absorbing capacity* 𝒜 : computed as *f*_tissue_ × *d*_structure_, the product of tissue area fraction *f*_tissue_ (foreground pixel fraction of the binarized mask) and median structure thickness *d*_structure_ (median Euclidean distance-transform value within the tissue phase) from the colony-surface ROIs, representing the total hygroscopic content per unit projected area. Specific surface area was computed as perimeter density (Sobel edge length per total area) divided by *f*_tissue_.

Colony-surface metrics (FFT spectral slope, thickness, absorbing capacity) and light-microscopy metrics (Hessian tubeness CV, erosion survival) were compared using Welch’s *t*-test as the primary test, with Mann–Whitney *U* as a secondary non-parametric test; for the light-microscopy comparison the exact Mann–Whitney *p*-value is bounded by *p* ≥ 0.024 at *n*_*a*_ = 6, *n*_*m*_ = 3 (limited by sample size). The four morphological metrics reduce to three underlying axes, volumetric tissue load (thickness, erosion survival), surface roughness (FFT spectral slope), and intra-tissue heterogeneity (Hessian tubeness CV), so the Bonferroni threshold at *α/*4 = 0.0125 conservatively accounts for the effective dimensionality of three; all four metrics exceed this threshold. Effect sizes are reported as Cohen’s *d*.

### Image analysis for field specimens

Field-collected *Gymnosporangium* specimens had imaging challenges that precluded automated Cellpose segmentation: low contrast between microdroplets and the heterogeneous leaf-surface background, inconsistent illumination across the curved leaf topography, and color variability from rust pigmentation and chlorophyll autofluorescence. Therefore, manual expert annotation was performed for all six field trials.

For each trial, the operator reviewed the complete time-lapse video sequence prior to annotation in order to establish temporal context: genuine condensation droplets grow and eventually shrink during evaporation, whereas static leaf-surface features do not exhibit morphological changes over time. Raw images were also pre-processed using ilastik (v1.4) [57] pixel classification as a contrast-enhancement tool to aid human visual perception of the images. All final boundary decisions were left to the human operator. Annotated boundaries were converted to binary masks and processed through the same morphometric pipeline used for laboratory data. Conservative inclusion criteria were applied to ensure reliable identification of condensation droplets: candidates were classified as droplets only if they exhibited growth over time followed by subsequent shrinkage due to evaporation. Any feature smaller than approximately 10 µm in diameter or directly adjacent to fungal hyphae was excluded from consideration as a droplet.

### Damköhler number estimation

The mass-transfer coefficient 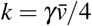 was determined based on 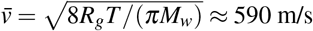 at 300 K [58]. For *L* = 1 mm and *D* = 2.5 × 10^−5^ m^2^/s, we find that *Da*_*s*_ ≈ 6 × 10^3^ for *γ* = 1 and *Da*_*s*_ ≈ 600 for *γ* = 0.1. These estimates confirm that uptake occurs under diffusion-limited conditions. This estimate uses the Hertz–Knudsen sticking coefficient for a flat liquid interface; for porous hydrogel or chitin surfaces the effective uptake kinetics may differ, but even a 10-fold further reduction in *γ* would yield *Da*_*s*_ ≈ 60, still within the diffusion-limited regime. However, since this calculation considers only the surface uptake step, additional considerations are needed for internal mass transfer. Transport of absorbed water into the bulk hyphal phase is governed by an internal effective diffusivity *D*_eff,wall_ ∼ 10^−10^ m^2^/s in chitin-rich walls [8], giving an internal mass-transfer coefficient *k*_int_ ∼ *D*_eff,wall_ / *d*_structure_ 5 × 10^−4^ m/s and an internal Damköhler number *Da*_*s*_ ∼ 1. It should be noted that the thin biological structures used here lie near the transition between diffusion-limited and mass-transfer-limited regimes, consistent with our use of a time-integrated effective-water-activity interpretation throughout this work.

### Statistical analysis

Ordinary least squares (OLS) regression was used to fit calibration curves on individual trials (*n* = 20; four hydrogel conditions). Cross-metric universal regressions (Fig. 3D,E) were performed using all 35 individual trials with unweighted OLS. Weighted OLS (weights = 1/SEM^2^) was applied for group-mean analyses. For the field validation, a sign test was used to assess whether all size–distance slopes were positive (*N* = 6). Pearson correlation was used for individual field trial regressions, with Spearman rank correlation as a nonparametric alternative where indicated. Bootstrap confidence intervals for calibration slopes were computed using *N* = 10,000 resamples (Supplementary Fig. 6). Leave-one-trial-out cross-validation of *δ* versus (1 − *a*_*w*_) yielded *R*^2^ = 0.896. Sample sizes were as follows: *n* = 5 per genus (15 fungi total), *n* = 5 per hydrogel condition (20 hydrogel trials total), *N* = 6 field trials. For colony architecture, *n* = 13 *Aspergillus* and *n* = 11 *Mucor* ROIs were used for colony-surface metrics, and *n* = 6 *Aspergillus* and *n* = 3 *Mucor* images for light-microscopy metrics. Shaded bands on profile plots (Figs. 2E, 3B, 5C) denote ± 1 s.e.m., while error bars on scatter plots and all text values denote ± 1 s.d. unless otherwise stated. All statistical analyses were carried out in Python 3.11 with SciPy, NumPy, and lifelines.

### Field collection

Samples of *Gymnosporangium* were collected in October 2025 from infected apple leaves on the Cornell University campus. Leaves bearing visible rust aecia were excised and transported to the laboratory the same day. A subset of these leaves had secondary fungal colonization and were classified as “diseased” specimens. Leaves were imaged under the same standardized condensation protocol, with the adaxial surface in direct contact with the cooled substrate.

## Data Availability

Raw time-lapse images and segmentation masks are deposited at Zenodo (https://doi.org/10.5281/zenodo.19391416).

## Code Availability

Analysis code and figure-generation scripts are available at https://github.com/Yany-Lin/fungal-condensation.

## Acknowledgements

This work was supported by the National Science Foundation under Grant No. CBET-2401507. We thank Zhengyang Liu and Yicong Fu for helpful discussions and technical assistance.

## Author Contributions

**Y.J.L**.: Conceptualization, Methodology, Software, Formal Analysis, Investigation, Visualization, Writing – Original Draft, Writing – Review & Editing. **L.F**.: Writing – Review & Editing. **A.K**.: Resources, Conceptualization, Writing – Review & Editing. **K.P**.: Funding Acquisition, Conceptualization, Writing – Review & Editing. **S.J**.: Conceptualization, Investigation, Supervision, Funding Acquisition, Writing – Review & Editing.

