## Supplementary Material for "Fungal Hyphae as Distributed Vapor Sinks"

### Supplementary Text 1: Internal equilibration and $a_{w,\mathrm{eq}}$ interpretation

The vapor penetration depth in chitin-rich walls is $\ell_{\mathrm{pen}} \sim \sqrt{D_{\mathrm{eff,wall}} \cdot t} \approx 300\ \mu\mathrm{m}$ over the 15-min condensation window, using a literature effective wall diffusivity $D_{\mathrm{eff,wall}} \sim 10^{-10}\ \mathrm{m^2/s}$ (Rosa et al. 2010; ref. 8 in the main paper), and exceeds either genus's structure thickness (4–7 $\mu$m) by more than an order of magnitude. The polymer phase therefore equilibrates internally within a second, so specific surface area is not rate-limiting. What matters instead is total absorbing mass per projected area: at the same external vapor flux, *Aspergillus*'s larger reservoir takes longer to fill, its surface water activity rises more slowly, and its time-averaged driving force $(1 - a_{w,\mathrm{eff}})$ stays larger across the experimental window, matching the $\sim 2\times$ ratio in measured $\delta$. Direct gravimetric tracking at colony resolution would test this mechanism directly but is beyond the scope of the present work.

The calibration returns effective water-activity values of $\sim 0.98$–$0.99$ for *Mucor* and *Rhizopus* (near the upper resolution limit of the calibration) and $\sim 0.93$ for *Aspergillus*. The thermodynamically relevant driving force $(1 - a_{w,\mathrm{eff}})$ therefore differs by roughly $3.5\times$ between *Aspergillus* and the other two genera (0.07 vs. 0.02). The dependence of $\delta$ on structure thickness implies non-equilibrium internal absorption during the 15-min window: the calibrated value is best interpreted as a time-integrated effective driving force rather than an instantaneous bulk water activity.

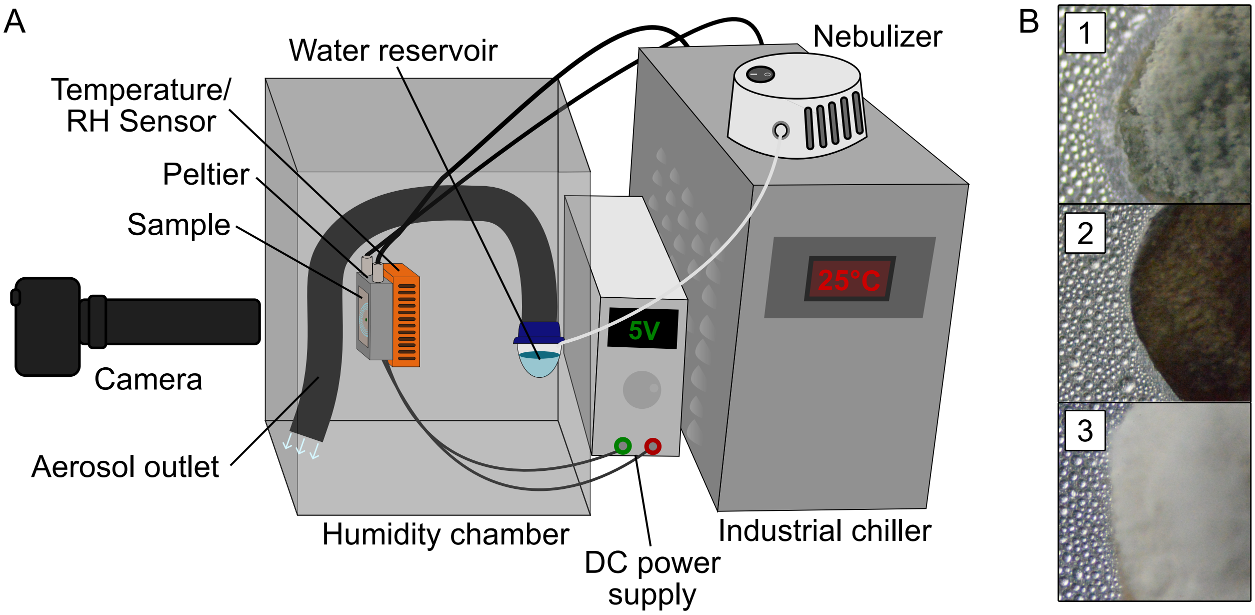


Supplementary Fig. 1: **Experimental setup and representative samples.** **(A)** Schematic of the sealed imaging chamber. A Peltier-cooled aluminum substrate ($T_s \approx 7\,^\circ$C) is thermally coupled to a closed-loop water chiller (CW-3000) via thermal paste. Silane-treated aluminum foil serves as the condensation surface. Water vapor is supplied by a compressor nebulizer (InnoSpire Essence, Respironics). Relative humidity and temperature are logged at 1-s resolution by a digital hygrometer–thermometer (IPT-100S, Elitech) with an external probe bound to the Peltier surface. Time-lapse imaging uses a DSLR camera on a motorized focus rail (StackShot, Cognisys) with LED illumination. **(B)** Representative brightfield images of samples on the cooled substrate: (1) a fungal colony patch, (2) a NaCl–agar hydrogel disk, and (3) an agar-only control. Circular patches are 3 mm in diameter.

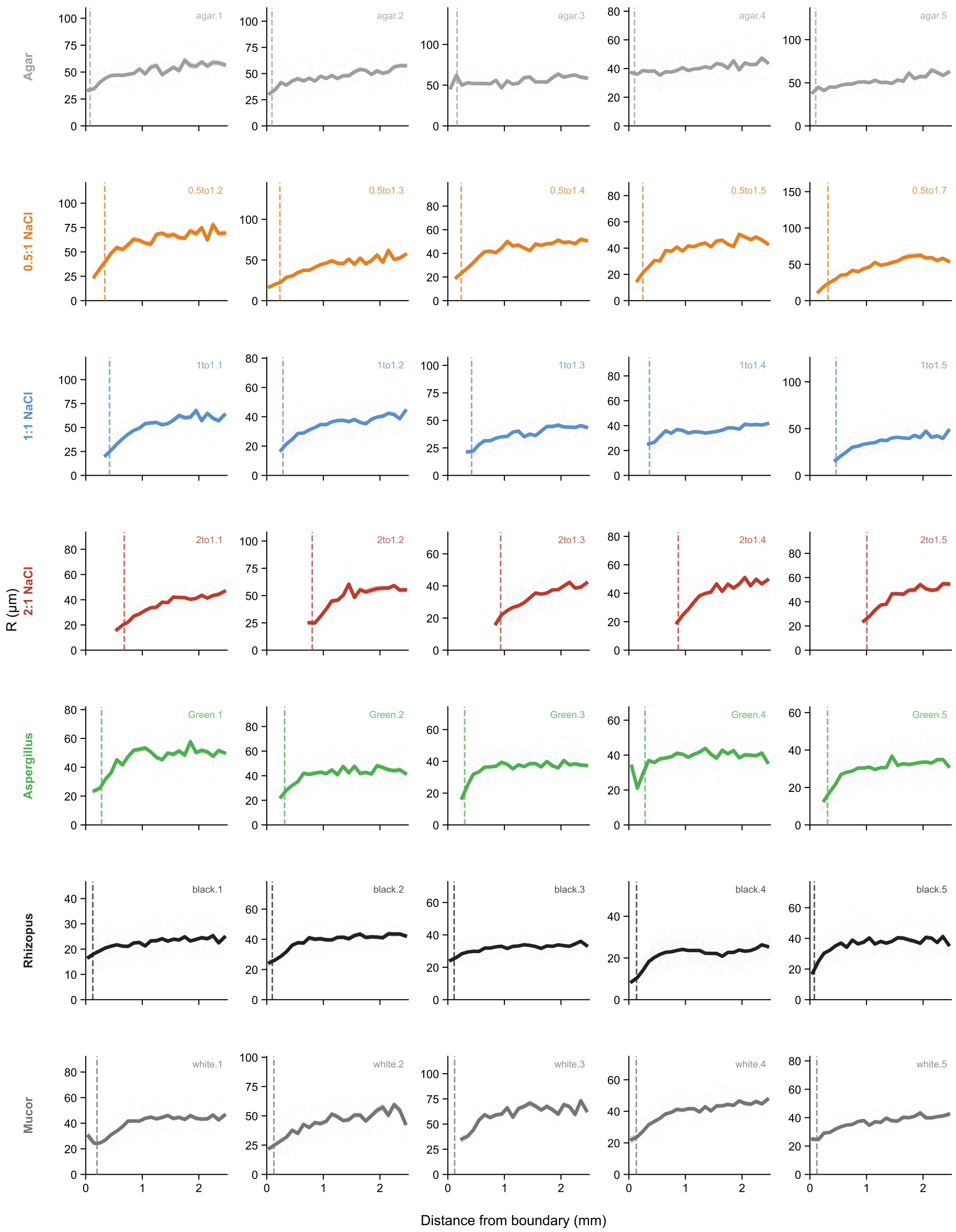


Supplementary Fig. 2: **Normalized radial size profiles for all 35 laboratory trials.** Each panel shows one trial (7 conditions × 5 replicates). Gray scatter: individual droplet radii at $t = 14.5$–$15.5$ min versus distance from the source boundary. Colored band: mean ± SEM computed in 100 $\mu$m bins. Red dashed line: dry-zone width $\delta$. Trial identifier is shown in the top-right corner of each panel. Conditions (top to bottom): agar, 0.5:1 NaCl, 1:1 NaCl, 2:1 NaCl, *Aspergillus*, *Rhizopus*, *Mucor*. The spatial size gradient—small near-field droplets transitioning to larger far-field droplets—is reproducible across all replicates and strengthens monotonically with increasing vapor-sink strength.

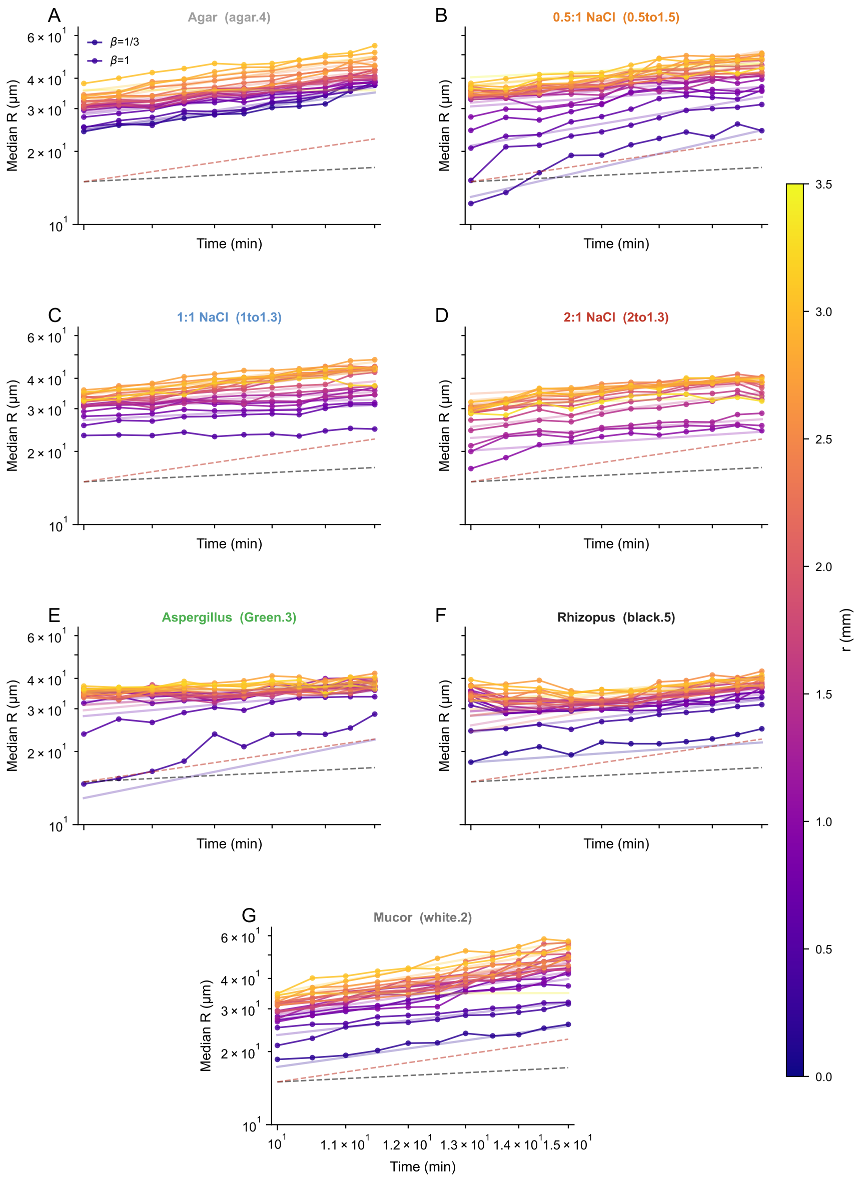


Supplementary Fig. 3: **Droplet growth curves for all seven conditions.** Each panel shows one representative trial per condition (**A**–**G**). Log–log plot of median droplet radius $R$ versus time, with droplets binned by distance from the source boundary (plasma colormap; navy = near, yellow = far). Solid colored lines: fitted power-law slopes. Dashed reference lines: diffusion-limited scaling $R \propto t^{1/3}$ ($\beta = 1/3$, black) and coalescence-dominated scaling $R \propto t^1$ ($\beta = 1$, red). Near-field droplets track the $t^{1/3}$ regime for longer across all conditions, consistent with growth suppression in the depletion zone delaying the transition to coalescence-dominated growth.

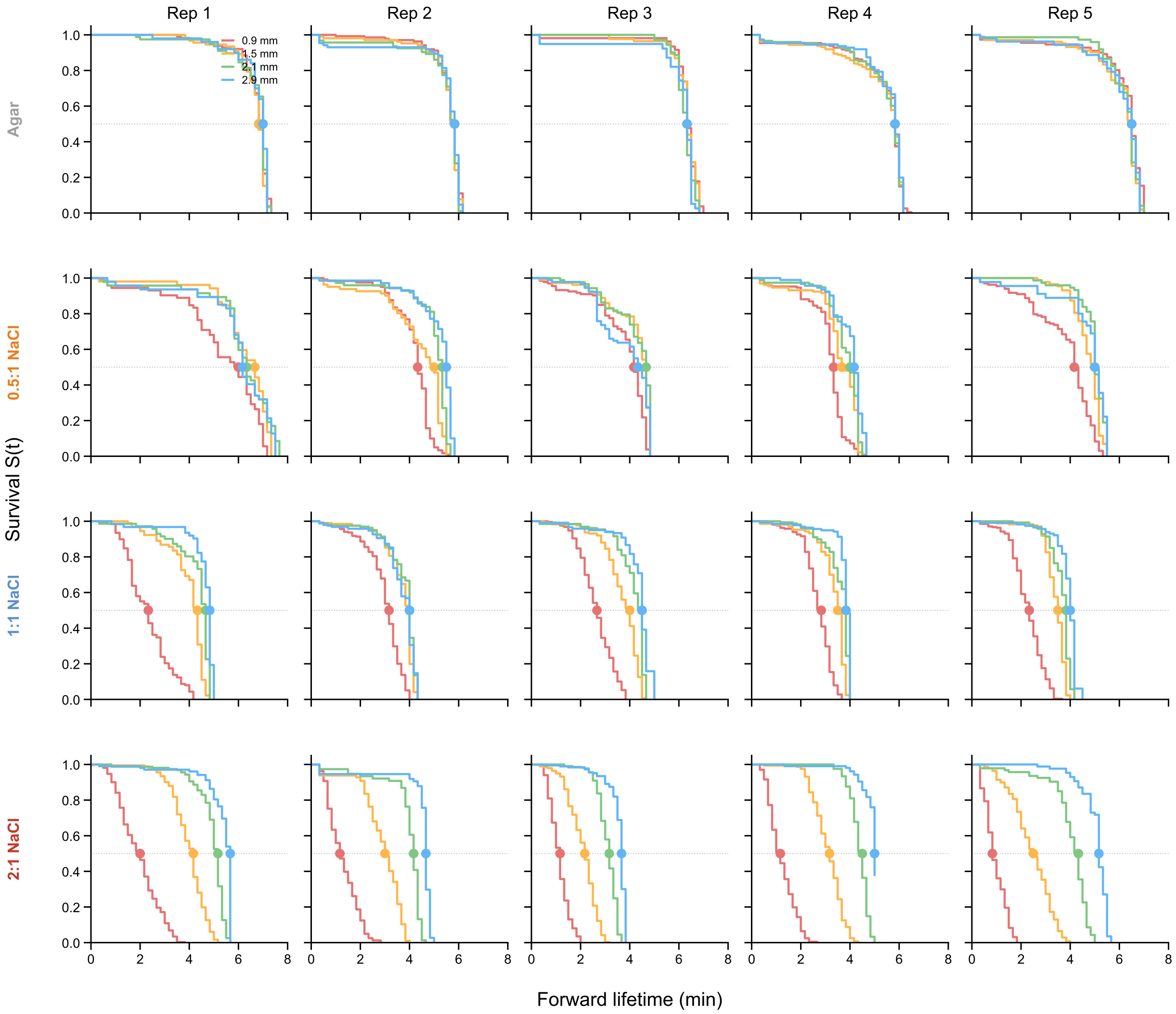


Supplementary Fig. 4: **Distance-stratified Kaplan–Meier survival curves for all 20 hydrogel trials.** Each panel shows one hydrogel trial (4 conditions $\times$ 5 replicates). Survival curves are stratified by four distance bands from the source boundary (0.9, 1.5, 2.1, and 2.9 mm). Dots mark the median survival time $\tau_{50}$ on the 0.5 survival line. Shaded bands: 95% confidence intervals. Forward lifetimes are measured from evaporation onset ($t = 15$ min). Agar controls show flat or weakly separated curves; 2:1 NaCl trials show the strongest distance-dependent survival gradient. The monotonic decrease in $\tau_{50}$ with proximity to the source is present in every replicate of every NaCl condition.

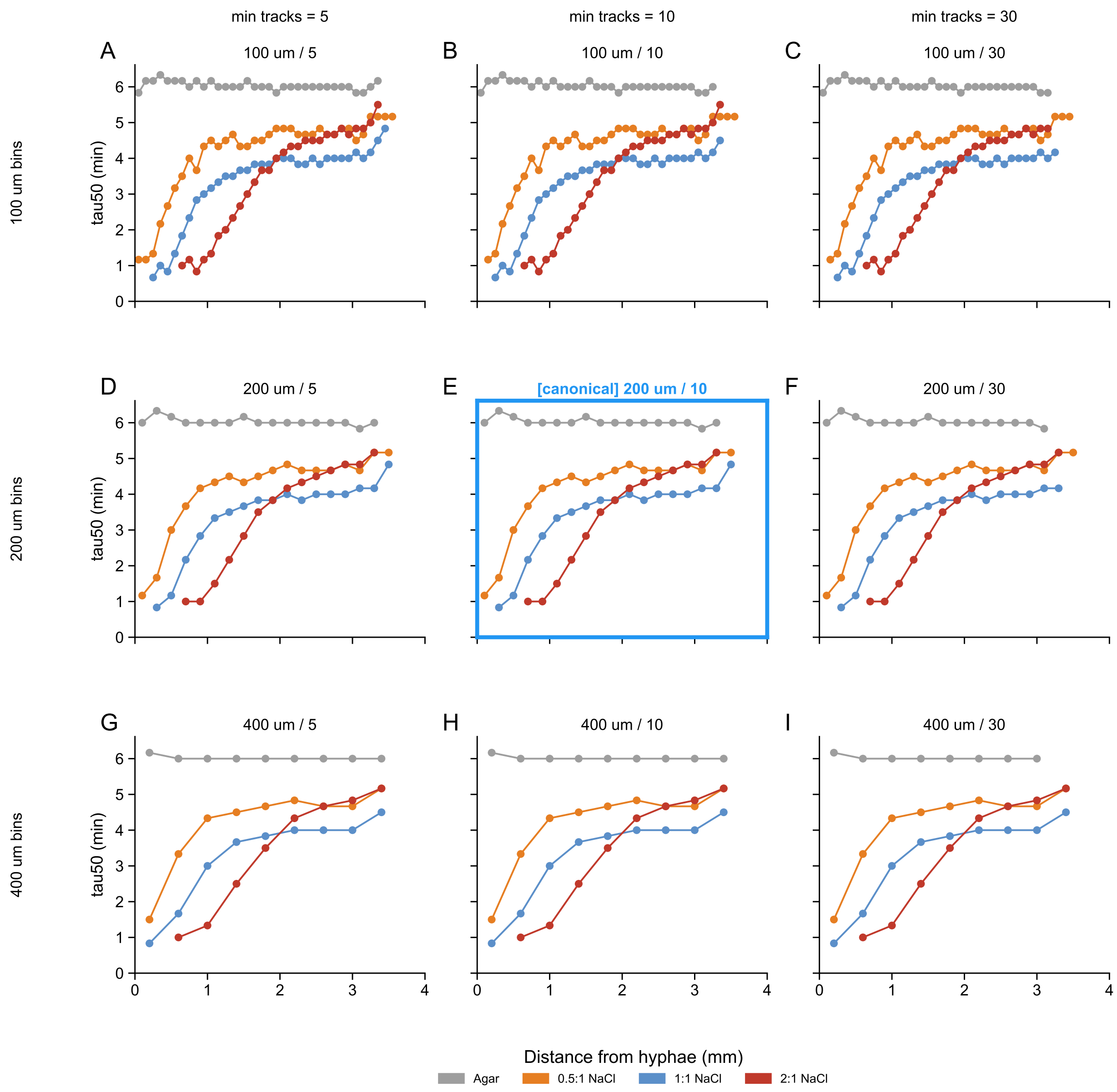


Supplementary Fig. 5: **Sensitivity of $\tau_{50}$–distance profiles to binning parameters.** $3 \times 3$ grid showing $\tau_{50}$ versus distance profiles for all four hydrogel conditions under nine combinations of bin width (100, 200, 400 $\mu$m) and minimum track count per bin (5, 10, 30). The canonical parameter set used in the main analysis (bin width = 200 $\mu$m, minimum tracks = 10; blue border) produces $\tau_{50}$ estimates that are consistent with alternative parameter choices. Profile shape and monotonic distance dependence are preserved across all nine combinations, confirming that the survival gradient metric is not sensitive to binning choices.

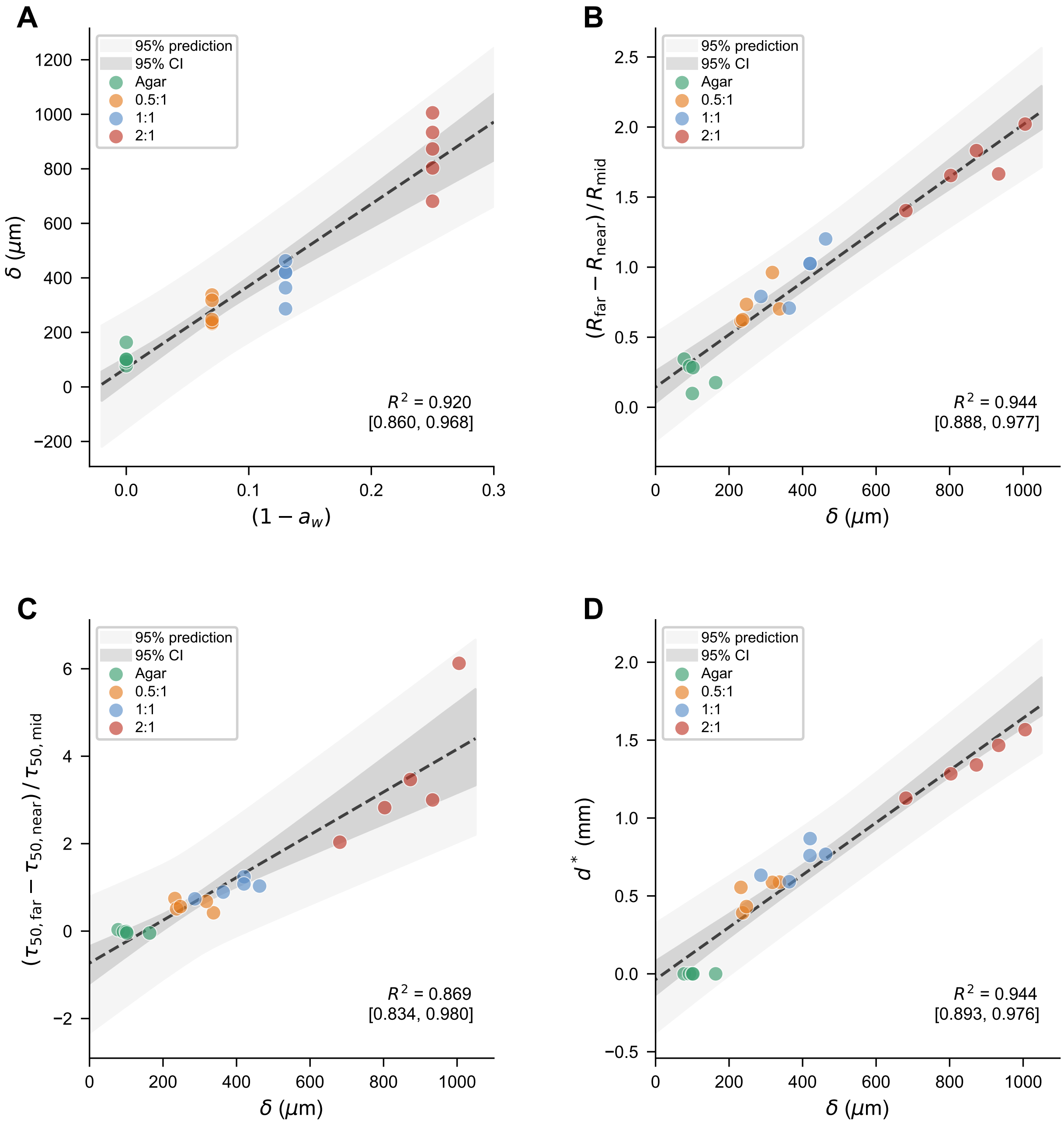


Supplementary Fig. 6: **Bootstrap confidence intervals for calibration regressions.** $2 \times 2$ panel grid showing the four calibration regressions from Fig. 2 with bootstrap 95% confidence and prediction bands ($N = 10{,}000$ resamples). **(A)** Dry-zone width $\delta$ versus $(1 - a_w)$ (Fig. 2F). **(B)** Normalized size gradient $(R_{\text{far}} - R_{\text{near}})/R_{\text{mid}}$ versus $\delta$ (Fig. 2H). **(C)** Normalized survival gradient $(\tau_{50,\text{far}} - \tau_{50,\text{near}})/\tau_{50,\text{mid}}$ versus $\delta$ (Fig. 2K). **(D)** Half-attenuation distance $d^*$ versus $\delta$ (Fig. 2L). Dark shading: 95% CI on the regression line; light shading: 95% prediction interval. Points are colored by hydrogel group (agar, 0.5:1, 1:1, 2:1 NaCl). Inset $R^2$ values with bootstrap 95% CIs confirm that all four linear relationships are statistically robust.

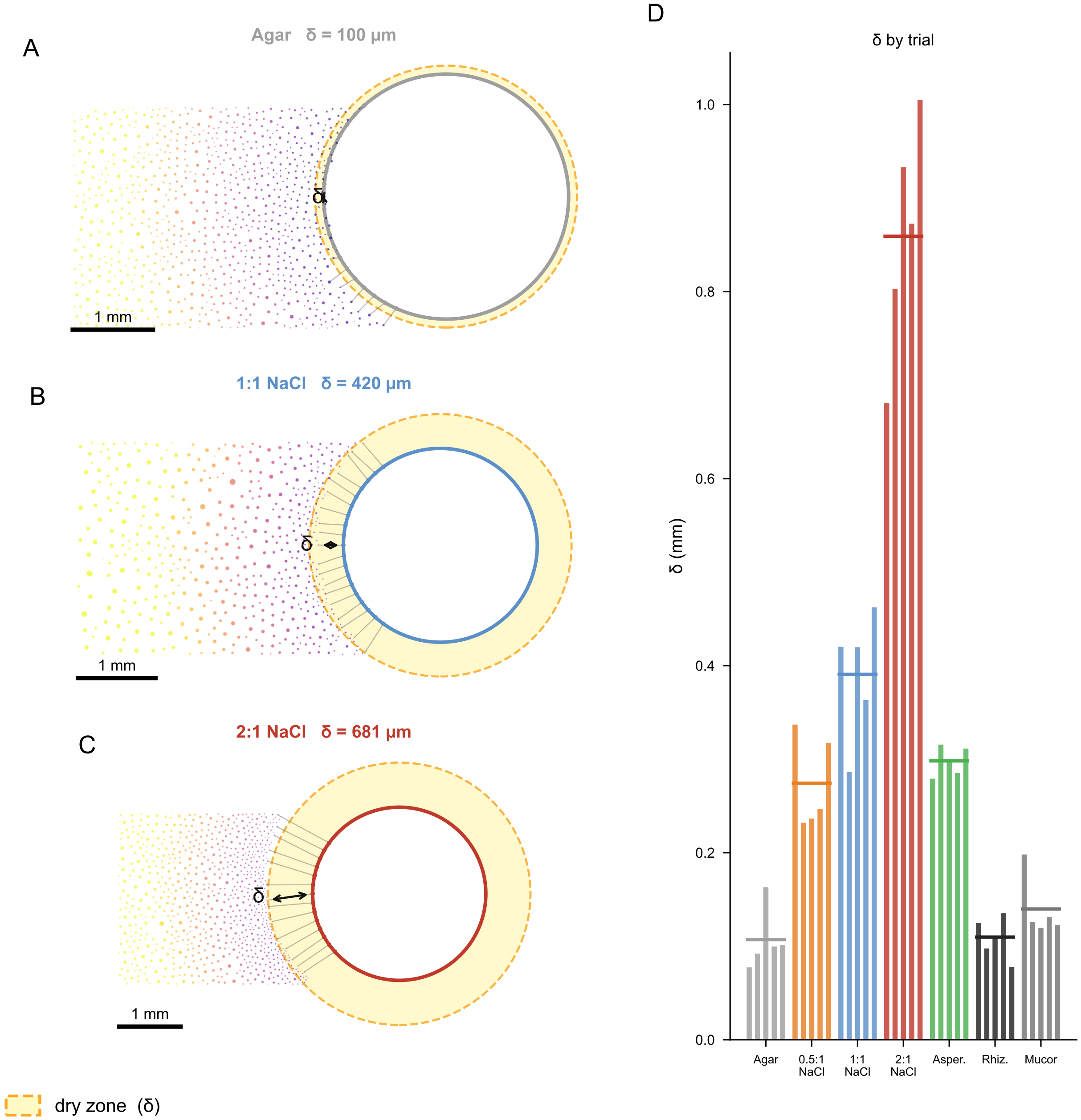


Supplementary Fig. 7: **Dry-zone width measurement by boundary raycast. (A–C)** Exemplar trials at three vapor-sink strengths: agar control ($\delta \approx 100\ \mu$m), 1:1 NaCl ($\delta \approx 420\ \mu$m), and 2:1 NaCl ($\delta \approx 681\ \mu$m). Each panel shows the droplet scatter colored by distance from the source boundary, the traced boundary polygon, and raycast lines used to sample the minimum droplet-free distance along 100 equally spaced boundary points. **(D)** Bar chart of $\delta$ for all 35 laboratory trials, grouped by condition. $\delta$ increases monotonically with vapor-sink strength across both hydrogel and fungal conditions.

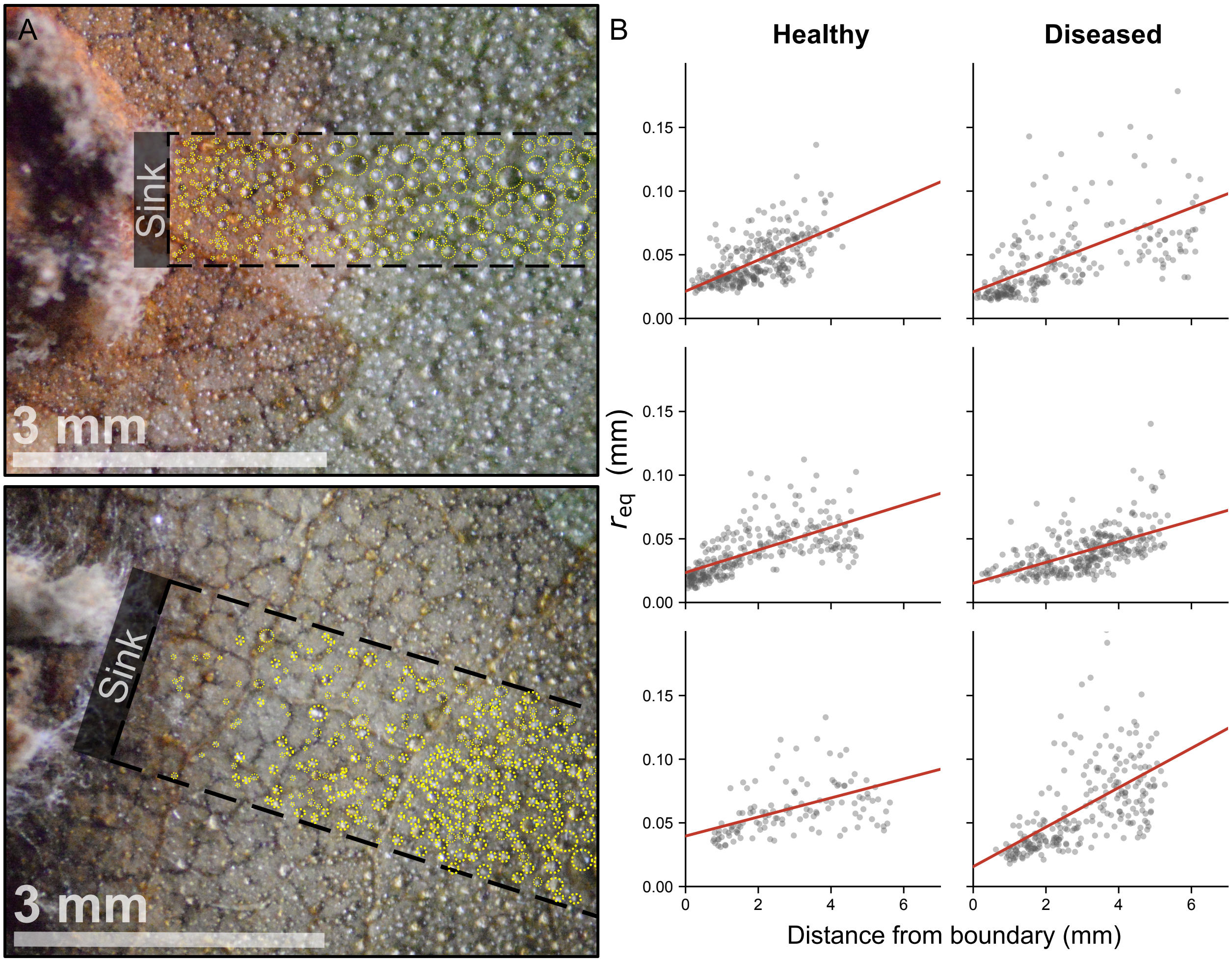


Supplementary Fig. 8: **Individual field trial size–distance scatter plots.** **(A)** Condensation micrographs of two *Gymnosporangium*-infected *Malus* leaf specimens with manual droplet annotations (red circles) and analysis regions (dashed rectangles). Top: healthy rust specimen. Bottom: diseased specimen with secondary fungal colonization. Scale bars: 3 mm. **(B)** Equivalent radius $r_{eq}$ versus distance from the fungal boundary for each of the six field trials (3 healthy, left column; 3 diseased, right column). Gray dots: individual annotated droplets ($N = 1{,}507$ total). Red lines: per-trial ordinary least-squares regression used to compute the size–distance slope reported in the main text. All six slopes are positive (Pearson $r = 0.43$–$0.66$, all $p < 10^{-6}$; sign test for all-positive slopes: two-tailed $p = 0.031$).

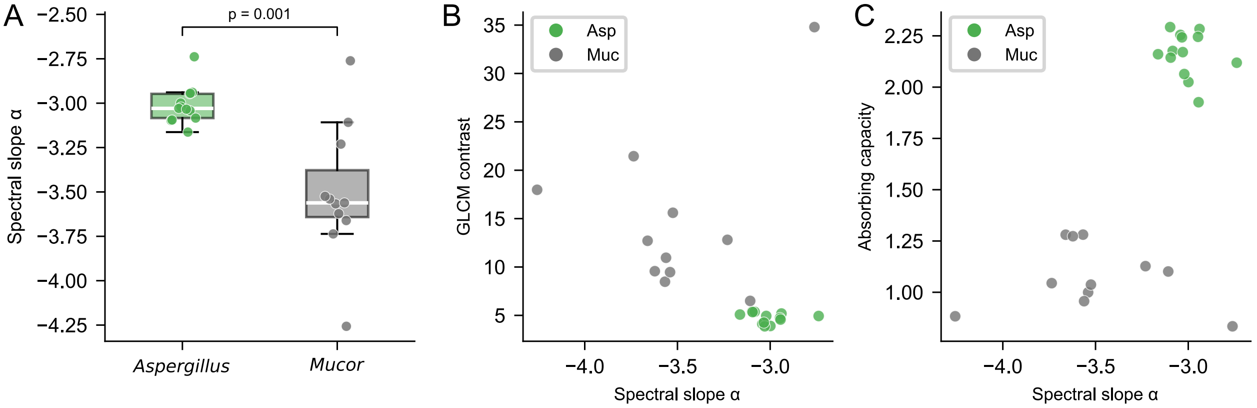


Supplementary Fig. 9: **FFT spectral slope and cross-validation.** **(A)** Tile-based 2D radial-PSD spectral slope $\alpha$ ($P(f) \propto f^{-\alpha}$) from macro-scale brightfield colony surfaces. *Aspergillus* ($-3.01 \pm 0.11$, $n = 13$) is shallower than *Mucor* ($-3.51 \pm 0.38$, $n = 11$; Welch's $t$-test, $p = 0.001$), reflecting greater sub-resolution surface roughness. **(B)** Cross-validation against an independent spatial-domain texture metric (GLCM contrast). **(C)** Spectral slope $\alpha$ versus absorbing capacity, confirming that surface texture covaries with the architectural mass-per-footprint metric. Scatter points in (B) and (C) color-coded by genus (*Aspergillus* green, *Mucor* gray). Boxes (A): IQR; whiskers: 1.5×IQR; line: median.

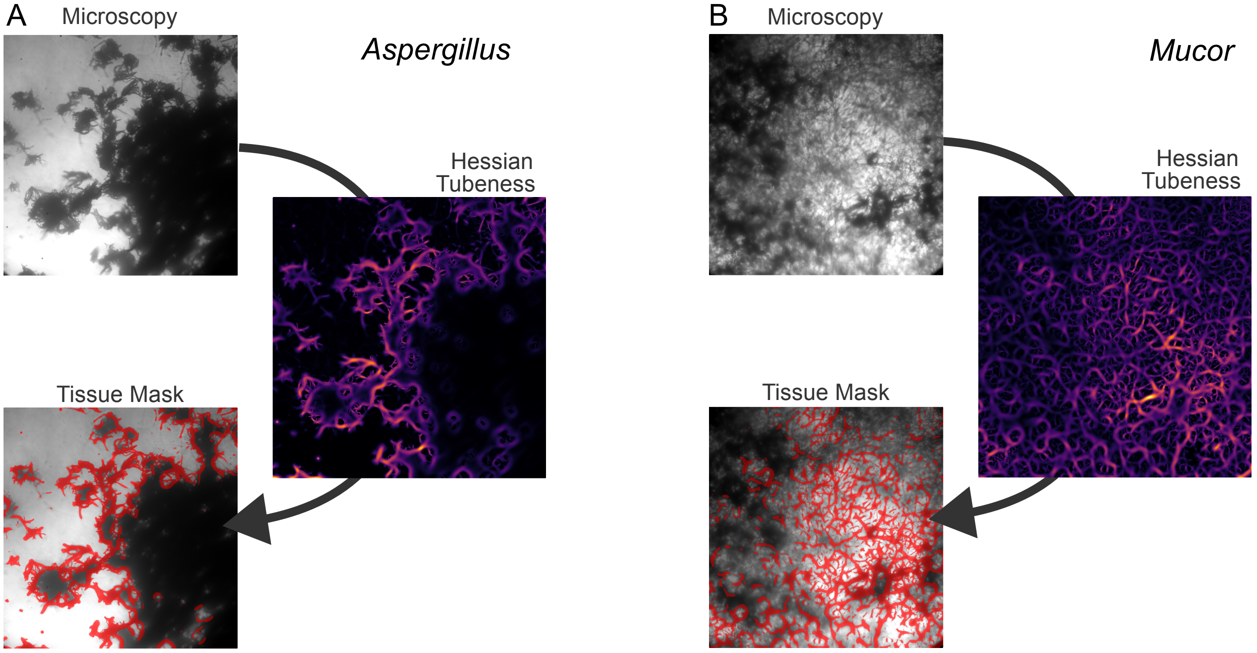


Supplementary Fig. 10: **Hessian tubeness pipeline on light microscopy.** Representative ROIs for *Aspergillus* **(A)** and *Mucor* **(B)**, showing the three-stage pipeline: original brightfield microscopy (10×), multi-scale Hessian tubeness response ($\sigma = 1$–$16$ px, max-projected; inferno colormap), and the corresponding adaptive-threshold tissue mask (red overlay). *Aspergillus* shows broad solid cores plus filamentous edges (high tubeness CV); *Mucor* is more uniformly filamentous (low tubeness CV).

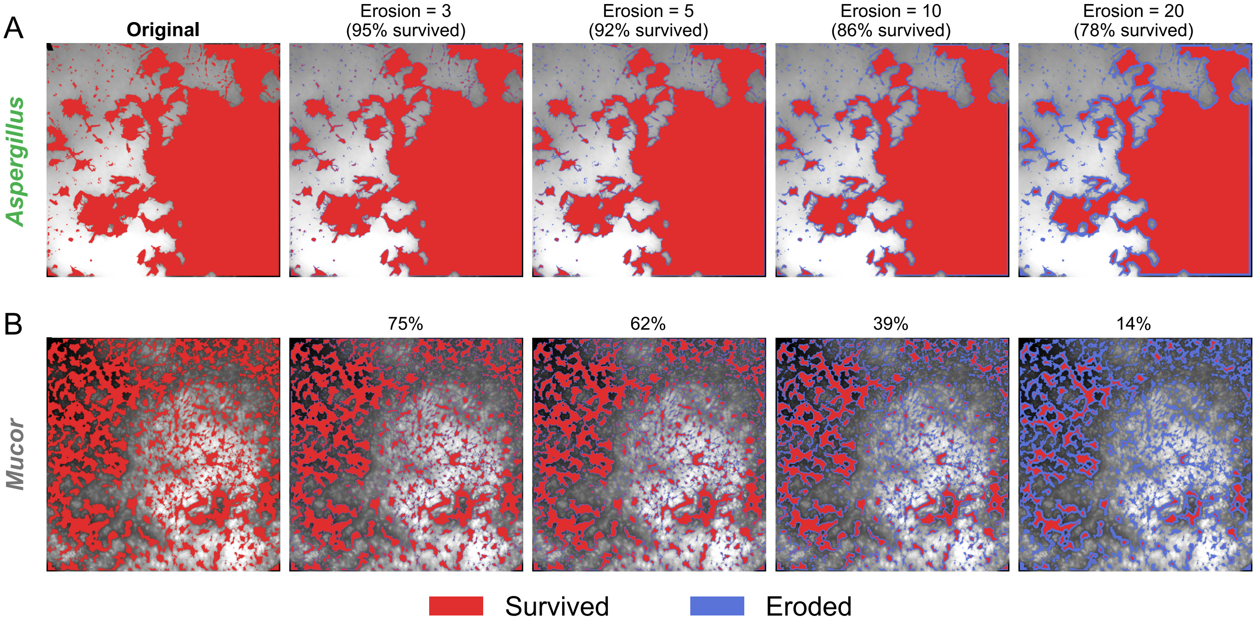


Supplementary Fig. 11: **Iterative erosion of light-microscopy tissue masks.** Representative *Aspergillus* **(A)** and *Mucor* **(B)** masks shown at iterations 0 (original), 3, 5, 10, and 20 (1-pixel-radius disk per iteration). Surviving tissue (red) and pixels removed at the current iteration (blue) are overlaid; percent retention is annotated above each frame. *Aspergillus* retains 86% at iteration 10 in this representative ROI (group mean 0.791; Fig. 4F); *Mucor* retains 39% (group mean 0.372). The disparity indicates thinner, more easily eroded filamentous structures in *Mucor*.

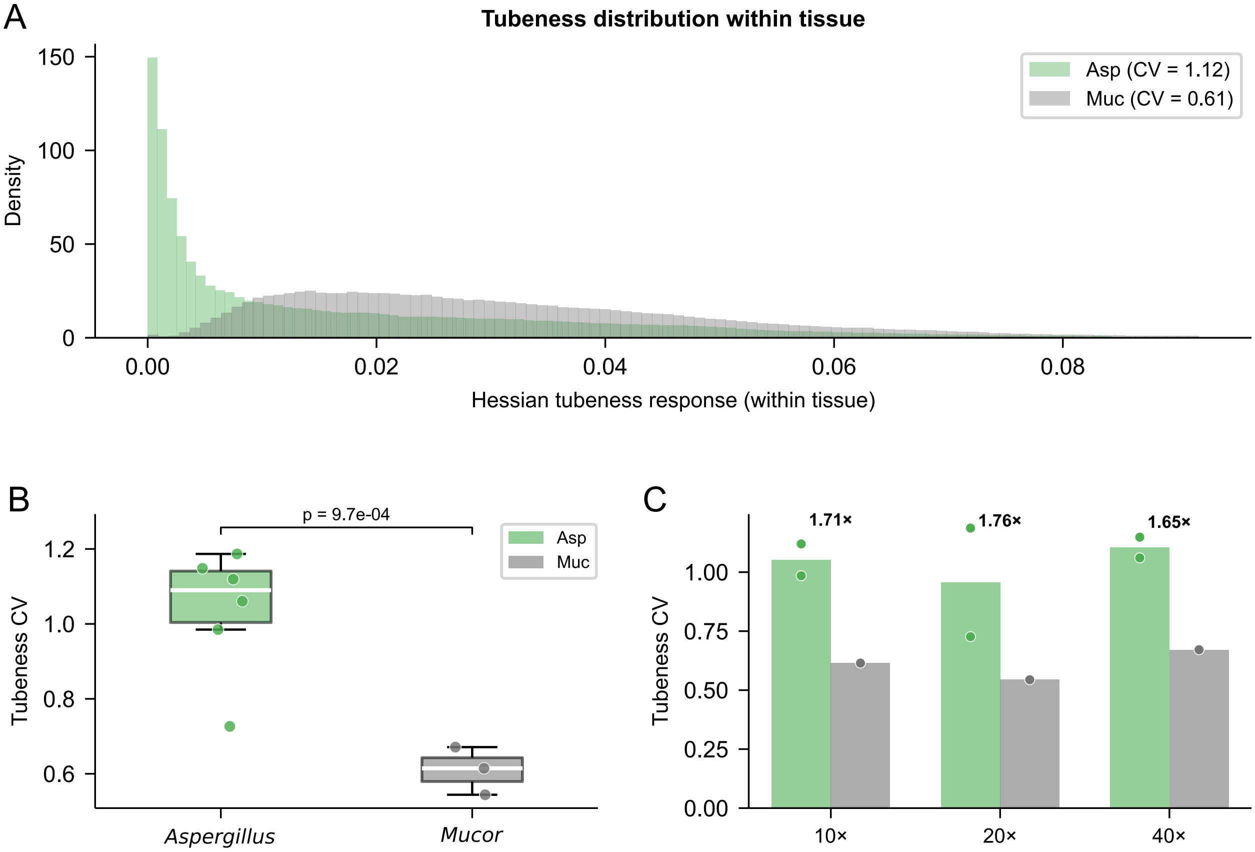


Supplementary Fig. 12: **Hessian tubeness CV: derivation and magnification stability.** **(A)** Within-tissue distribution of Hessian tubeness response for one representative ROI per genus. *Aspergillus* (green) is bimodal (low-tubeness solid cores plus high-tubeness filamentous edges, CV = 1.12); *Mucor* (gray) is more unimodal (CV = 0.61). **(B)** Per-image tubeness CV; Welch's $t$-test, $p = 9.7 \times 10^{-4}$ ($n_a = 6$, $n_m = 3$). **(C)** Asp/Muc tubeness CV ratio is preserved across magnifications: 1.71× at 10×, 1.76× at 20×, 1.65× at 40×. Boxes (B): IQR; whiskers: 1.5×IQR; line: median.

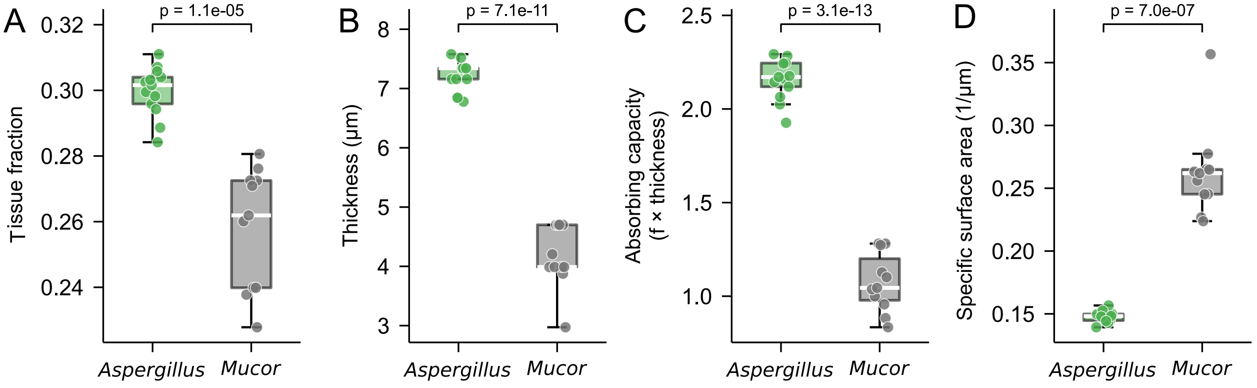


Supplementary Fig. 13: **Absorbing capacity decomposition.** Per-ROI box plots ($n = 13$ *Aspergillus*, $n = 11$ *Mucor*). **(A)** Tissue area fraction $f_{\text{tissue}}$ (Welch's $t$-test, $p = 1.1 \times 10^{-5}$). **(B)** Median structure thickness $d_{\text{structure}}$ ($p = 7.1 \times 10^{-11}$). **(C)** Absorbing capacity $\mathscr{A} = f_{\text{tissue}} \times d_{\text{structure}}$ ($p = 3.1 \times 10^{-13}$). **(D)** Specific surface area (perimeter / tissue area). *Mucor* exhibits 1.78× higher SSA than *Aspergillus* ($p = 7.0 \times 10^{-7}$); this inversion relative to vapor-sink strength indicates that surface-to-volume ratio is not the quantity controlling sink strength, and instead supports the absorbing-capacity (mass-per-footprint) interpretation. Boxes: IQR; line: median; whiskers: 1.5×IQR; dots: individual ROIs.

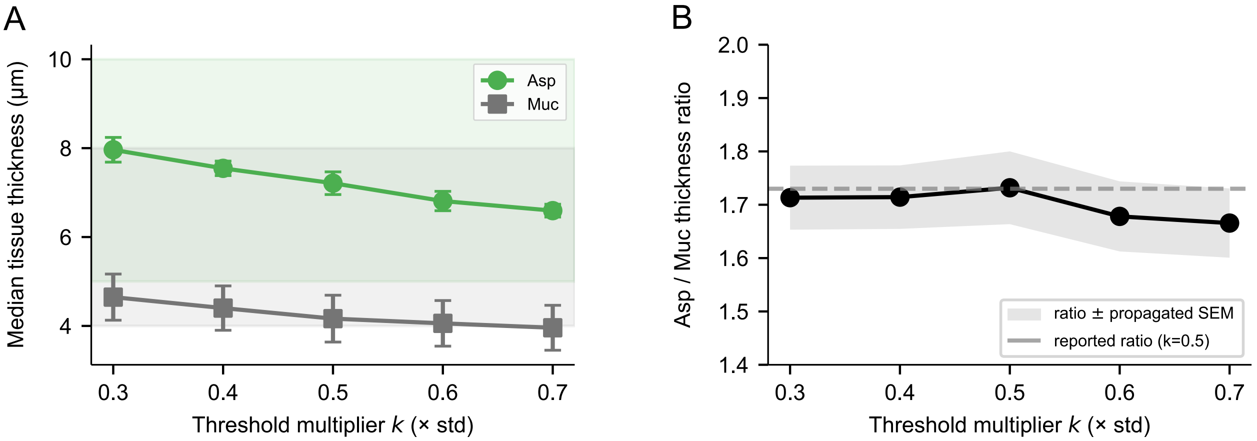


Supplementary Fig. 14: **Structure-thickness segmentation is robust to threshold choice.** **(A)** Median tissue thickness for *Aspergillus* (green) and *Mucor* (gray) as the adaptive threshold multiplier $k$ is swept from 0.3 to 0.7 × std. Both genera fall within published hyphal-diameter ranges (shaded bands) across the entire sweep. **(B)** Asp/Muc thickness ratio across the $k$ sweep. The ratio is robust to threshold choice (range 1.67–1.73), confirming that the headline 1.73× value reported at $k = 0.5$ (dashed line) is not a parameter-selection artifact. Bands: ± SEM (propagated from per-genus SEM).

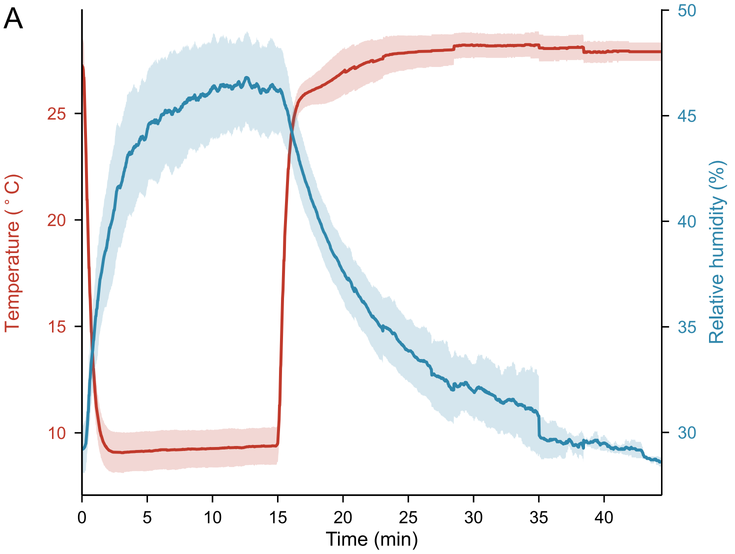


B

| **Metric** | **Temperature** | **Humidity** |
|---|---|---|
| n trials | 13 | 13 |
| Trajectory range | 9.1–28.2 °C | 28.6–46.8 % |
| Mean across-trial SD | 0.78 °C | 1.61 % |
| **Time-pooled SD (RMS)** | **0.81 °C** | **1.79 %** |

Supplementary Fig. 15: **Chamber temperature and humidity reproducibility across 13 trials. (A)** Mean trajectories of substrate temperature (red, left axis) and chamber relative humidity (blue, right axis) with $\pm 1$ SD bands across $n = 13$ standardized trials; condensation phase $t = 0$–15 min, evaporation $t > 15$ min. **(B)** Stability statistics: trajectory range, mean across-trial SD (0.78 °C, 1.61% RH), and time-pooled SD (root-mean-square of the across-trial SD time series; 0.81 °C, 1.79% RH) for both channels.

Supplementary Table 1: **Sensitivity of the normalized size gradient to zone boundaries.** The size gradient $(\bar{R}_{\text{far}} - \bar{R}_{\text{near}})/\bar{R}_{\text{mid}}$ was regressed against $\delta$ across all 35 laboratory trials for 36 combinations of near, mid, and far evaluation zones. All combinations give $R^2 \geq 0.82$ and Spearman $\rho \geq 0.81$. The canonical zones (bold; near 0–500, mid 900–1100, far 1900–2100 $\mu$m) yield $R^2 = 0.905$ ($p = 1.9 \times 10^{-18}$, $n = 35$); this is the same regression reported as $R^2 = 0.91$ (rounded) in the main text.

| Near ($\mu$m) | Mid ($\mu$m) | Far ($\mu$m) | $n$ | $R^2$ | $\rho$ |
|---|---|---|---|---|---|
| 0–700 | 900–1100 | 1500–2500 | 35 | 0.928 | 0.852 |
| 0–700 | 900–1100 | 1700–2300 | 35 | 0.928 | 0.851 |
| 0–700 | 800–1200 | 1700–2300 | 35 | 0.927 | 0.852 |
| 0–700 | 800–1200 | 1500–2500 | 35 | 0.926 | 0.844 |
| 0–1000 | 750–1250 | 1700–2300 | 35 | 0.924 | 0.832 |
| 0–1000 | 750–1250 | 1500–2500 | 35 | 0.923 | 0.812 |
| 0–1000 | 800–1200 | 1700–2300 | 35 | 0.923 | 0.832 |
| 0–1000 | 800–1200 | 1500–2500 | 35 | 0.923 | 0.811 |
| 0–700 | 900–1100 | 1900–2100 | 35 | 0.922 | 0.851 |
| 0–700 | 800–1200 | 1900–2100 | 35 | 0.922 | 0.846 |
| 0–700 | 750–1250 | 1500–2500 | 35 | 0.920 | 0.835 |
| 0–700 | 750–1250 | 1700–2300 | 35 | 0.920 | 0.851 |
| 0–1000 | 750–1250 | 1900–2100 | 35 | 0.919 | 0.827 |
| 0–1000 | 900–1100 | 1500–2500 | 35 | 0.918 | 0.820 |
| 0–1000 | 900–1100 | 1700–2300 | 35 | 0.918 | 0.832 |
| 0–1000 | 800–1200 | 1900–2100 | 35 | 0.917 | 0.829 |
| 0–700 | 750–1250 | 1900–2100 | 35 | 0.917 | 0.852 |
| 0–500 | 900–1100 | 1700–2300 | 35 | 0.912 | 0.861 |
| 0–1000 | 900–1100 | 1900–2100 | 35 | 0.911 | 0.825 |
| 0–500 | 900–1100 | 1500–2500 | 35 | 0.911 | 0.856 |
| 0–500 | 800–1200 | 1700–2300 | 35 | 0.907 | 0.858 |
| 0–500 | 800–1200 | 1500–2500 | 35 | 0.905 | 0.847 |
| **0–500** | **900–1100** | **1900–2100** | **35** | **0.905** | **0.864** |
| 0–500 | 800–1200 | 1900–2100 | 35 | 0.901 | 0.854 |
| 0–500 | 750–1250 | 1700–2300 | 35 | 0.895 | 0.860 |
| 0–500 | 750–1250 | 1500–2500 | 35 | 0.893 | 0.856 |
| 0–500 | 750–1250 | 1900–2100 | 35 | 0.890 | 0.855 |
| 0–300 | 900–1100 | 1700–2300 | 35 | 0.860 | 0.861 |
| 0–300 | 900–1100 | 1500–2500 | 35 | 0.860 | 0.858 |
| 0–300 | 900–1100 | 1900–2100 | 35 | 0.854 | 0.864 |
| 0–300 | 800–1200 | 1700–2300 | 35 | 0.850 | 0.855 |
| 0–300 | 800–1200 | 1500–2500 | 35 | 0.849 | 0.855 |
| 0–300 | 800–1200 | 1900–2100 | 35 | 0.845 | 0.855 |
| 0–300 | 750–1250 | 1700–2300 | 35 | 0.831 | 0.863 |
| 0–300 | 750–1250 | 1500–2500 | 35 | 0.830 | 0.857 |
| 0–300 | 750–1250 | 1900–2100 | 35 | 0.827 | 0.859 |

Supplementary Table 2: **Per-metric architecture statistics for *Aspergillus* vs. *Mucor*.** Welch's $t$-test as primary, with Mann–Whitney $U$ used as non-parametric confirmation (light-microscopy MWU bounded by $p \geq 0.024$ at $n_a = 6$, $n_m = 3$; not tabulated). All four metrics survive Bonferroni correction at $\alpha/4 = 0.0125$ against the effective dimensionality of three (volumetric hyphal load, surface roughness, intra-hyphal heterogeneity). $n = 13$ *Aspergillus* and $n = 11$ *Mucor* ROIs for FFT slope and structure thickness; $n = 6$ *Aspergillus* and $n = 3$ *Mucor* for tubeness CV and erosion survival.

| Metric | *Aspergillus* | *Mucor* | Ratio | Welch $p$ | Cohen $d$ |
|---|---|---|---|---|---|
| FFT spectral slope $\alpha$ | $-3.01 \pm 0.11$ | $-3.51 \pm 0.38$ | 1.17 ($\lvert\alpha\rvert$) | $1.0 \times 10^{-3}$ | 1.85 |
| Structure thickness ($\mu$m) | $7.21 \pm 0.26$ | $4.16 \pm 0.53$ | 1.73 | $7.1 \times 10^{-11}$ | 7.58 |
| Hessian tubeness CV | $1.04 \pm 0.17$ | $0.61 \pm 0.06$ | 1.70 | $9.7 \times 10^{-4}$ | 2.93 |
| Erosion survival (iter. 10) | $0.791 \pm 0.103$ | $0.372 \pm 0.025$ | 2.13 | $7.9 \times 10^{-5}$ | 4.75 |

### Supplementary Movie Legends

**Supplementary Movie 1. Time-lapse condensation around an agar control disk.** DSLR macro time-lapse of the condensation–evaporation cycle around a 3 mm agar-only disk ($a_w = 1.00$) on the cooled aluminum substrate. The non-hygroscopic control shows uniform droplet growth across the field with no near-source depletion zone, providing the no-sink baseline against which all hygroscopic samples are compared.

**Supplementary Movie 2. Time-lapse condensation around a 2:1 NaCl–agar hydrogel.** DSLR macro time-lapse around a 3 mm 2:1 NaCl–agar hydrogel disk ($a_w = 0.75$), the strongest abiotic sink in the calibration series. A clear dry zone forms within the first minute, near-source droplet growth is suppressed throughout condensation, and near-source droplets disappear first when evaporation begins.

**Supplementary Movie 3. Time-lapse condensation around an *Aspergillus* colony.** DSLR macro time-lapse around a 3 mm *Aspergillus* colony patch transferred onto the cooled substrate with aerial mycelium upward. The dense conidia-dominated surface produces the widest depletion zone of the three fungal genera, with near-source growth suppression and outward-propagating evaporation matching the laboratory hydrogel pattern.

**Supplementary Movie 4. Time-lapse condensation around a *Mucor* colony.** DSLR macro time-lapse around a 3 mm *Mucor* colony patch under matched conditions. The hypha-dominated surface produces a narrower depletion zone than *Aspergillus* despite higher chitin–chitosan content per unit mass, illustrating that absorbing capacity (mass per footprint), not surface-to-volume ratio, sets vapor-sink strength.

**Supplementary Movie 5. Time-lapse condensation on a *Gymnosporangium*-infected *Malus* leaf.** DSLR macro time-lapse of a representative healthy rust-infected *Malus* leaf imaged under the laboratory protocol with the adaxial surface in contact with the cooled substrate. Droplets near the orange-yellow rust aecium are smaller and disappear first as the evaporation front propagates outward, reproducing the laboratory vapor-sink signature in a natural pathosystem.
